# HIF1α gates tendon response to overload and drives tendinopathy independently of vascular recruitment

**DOI:** 10.1101/2025.02.02.635939

**Authors:** Greta Moschini, Archana G. Mohanan, Izabella S. Niewczas, Diane E. Taylor, Patrick K. Jaeger, Amro A. Hussien, Stefania L. Wunderli, Olivia Baumberger, Maja Wolleb, Barbara Niederoest, Maja Bollhalder, Raphaela Ardicoglu, Guillermo Turiel, Evi Masschelein, Sarah Morice, Santiago Ardiles, Lieke Mous, Matthew R. Aronoff, Monika Hilbe, Farah Selman, Karl Wieser, Sandro F. Fucentese, Fabian S. Passini, Ulrich Blache, Didier Surdez, Helma Wennemers, Dirk Elewaut, Jonathan Clark, Katrien De Bock, Jess G. Snedeker

## Abstract

Tendons are mostly avascular dense connective tissues that link muscles to bones, withstanding some of the highest mechanical stresses in the body. Mechanical overload and tissue hypervascularity are implicated in tendinopathy, a common musculoskeletal disorder, but mechanistic understanding of their roles is largely lacking. Here, we identify HIF1α not only as a marker but as a driver of tendinopathy. Initial histological and multi-omics evaluation of human tendinopathic samples revealed extensive extracellular matrix remodeling, including pathological collagen crosslinking coinciding with active hypoxic signaling. Hypothesizing a causal contribution of hypoxia signaling, we generated mice with tenocyte-targeted deletions of the Von Hippel-Lindau (VHL) gene, which controls hypoxia signaling by regulating HIFα degradation. We demonstrated that VHL inactivation suffices to induce pathological hallmarks of tendinopathy, such as collagen matrix disorganization, crosslinking, altered mechanics and neuro-vascular ingrowth. This phenotype was HIF1α-dependent, since co-deleting HIF1α rescued tendon morphology and mechanics. Moreover, deleting vascular endothelial growth factor A (VEGFA) alongside VHL effectively decoupled the effects of vascular ingrowth from persistently aberrant extracellular matrix remodeling and mechanical dysfunction, emphasizing a direct role of HIF1α in driving tendon disease that is independent of angiogenesis. Mechanistically, we linked tendon mechanical overload to the onset of HIF1α signaling in primary cultured human tendon cells. Furthermore, genetically removing HIF1α from tenocytes prevented aberrant tendon remodeling in response to chronic overload. These findings position HIF1α signaling as a central driver of tendinopathy that acts through a maladaptive tissue response to chronic overload, providing mechanistic insights that could be leveraged for improved therapeutic approaches.

**One Sentence Summary:** HIF1α activation promotes tendinopathy and its inhibition prevents overload-induced maladaptation, suggesting therapeutic potential.

## INTRODUCTION

By connecting muscle to bone, tendons play an essential role in human movement (^1^). Healthy tendons feature a tightly packaged type-I collagen extracellular matrix (ECM) that is extremely dense, contains relatively few tendon cells, is largely avascular and has a low state of metabolic activity (^2–5^). In contrast, diseased tendons are characterized by a loss of its tightly organized ECM, fibrosis, and increased presence of blood vessels, immune cells and nerves (^6–11^). Even though tendinopathy affects 2-5% of the general population at any given moment (^12^) and up to 22% of athletic population (^13^), it remains poorly understood and poses a significant socioeconomic burden (^14,15^).

The complex and unclear etiology of tendinopathy has led to several hypotheses about its pathogenesis with differing contribution of blood vessels (^2,11,16–19^). The mechanical hypothesis emphasizes the role of excessive mechanical loading that exceeds the intrinsic capacity of the tendon to adapt. This overload can result from repetitively high loads or increased exercise frequency, leading to tissue damage and failed healing (^1^). Under those conditions, the presence of blood vessels is the consequence of a failed healing response (^20^). Alternatively, the vascular hypothesis identifies pathological vascularity as an important driver of the disease, whereby infiltrating blood vessels cause matrix derangement, compromise functional load-bearing capacity of the organ and promote the ingrowth of pain-associated nerves, suggestive of a chronically active healing response (^21–23^).

Both theories have been indirectly related to hypoxia inducible factor 1α (HIF1α), a key transcriptional effector of the cellular response to hypoxia (^22^). HIF1α- dependent genetic programs promote cell survival under hypoxic conditions through modulation of metabolic pathways, increase tissue vascularity by increasing the expression of angiogenic growth factors, control proliferation and differentiation in many organs and are often associated with fibrosis (^24^). In the context of tendon pathology, increased HIF1α expression has been observed in diseased rotator cuff tendons (^25–29^) and tissue hypoxia has long been suspected to play a central role in progression of tendon disease (^30,31^). However, little mechanistic work has been focused on elucidating the upstream regulators, downstream effectors and molecular mechanisms that may link HIF1α to tendinopathy. Furthermore, it is unclear whether HIFs are sufficient and/or required for the development of tendinopathy.

In this study, we used integrative multi-omics analysis of human tendinopathic samples alongside genetic mouse models and *in vitro*/*in vivo* overload models to demonstrate that chronic HIF1α activation is a key driver of tendon pathology, driving tissue derangement in a manner independent of vascular ingrowth. Moreover, we show HIF1α activation in response to mechanical overload. Furthermore, inhibition of HIF1α prevented tendon maladaptation to overload, revealing it as a molecular target for potential therapeutic intervention against tendinopathy.

## RESULTS

### Diseased human tendons show substantially increased trivalent collagen crosslinking among other aberrant ECM changes

To investigate the molecular mechanisms underlying human tendinopathy, we selected and characterized a cohort of long head biceps tendons (LHBT) harvested from pathological shoulder joints (**Table S1**). Previous studies observed a higher degree of degeneration in the proximal LHBT region (proximal LHBT) compared to the corresponding distal region (distal LHBT) (^32–34^), we therefore analysed and compared these zones. In addition, an independent cohort of gracilis tendons was used as healthy control to assess the suitability of the distal LHBT as an intra-patient control (**Fig. 1A**). When comparing the overall histopathological grade (Bonar score) (^35^) among the three groups, we found that the proximal LHBT showed an advanced pathological stage compared to the distal LHBT (p=0.0068) and the gracilis tendon (p<0.0001) (**Fig. 1B**). In line with the Bonar score, immuno-histological characterization of the proximal LHBT samples displayed increased cell roundness, loss of matrix integrity, blood vessel infiltration (CD31^+^ cells) and ingrowth of myelinated nerve cells (NM-F+ cells) to the tendon proper. The cell number was comparable between the three groups (**Fig. 1C-D and fig. S1A**). Since matrix dysregulation is a central hallmark of tendon disease (^36^), we next performed quantitative proteomics of the most abundant collagen components (^37^). We found reduced COL1A1 and COL2A1 protein levels in the proximal LHBT compared to the distal LHBT and gracilis controls. A similar trend was observed for COL4. By contrast, COL3A1 and COL5 protein expressions were highly increased (p<0.0001) in the proximal LHBT area (**Fig. 1E and fig. S1B**). We then determined whether these changes in collagen content were accompanied by altered collagen crosslinking, which is typically observed in fibrosis (^38^). Using mass-spectrometry (MS), we found a 3-fold increase in the mature crosslinks pyridinoline (PYD) and deoxy-pyridinoline (DPD) in the proximal LHBT region compared to distal LHBT and gracilis controls. A similar increase was observed in the proximal LHBT for the immature crosslinks dihydroxylysino-norleucine (DHLNL) and hydroxylysino-norleucine (HLNL), while the total glycation levels (Hly+Lys) were lower compared to the other two groups (**Fig. 1F and fig. S1C**). These observations are consistent with higher rates of collagen turnover (^39^). Taken together, these results indicate that proximal LHBT samples compared to the distal LHBT region and the healthy gracilis tendons feature sharply distinct collagen crosslink characteristics, among other more recognized features of tendinopathy, including ECM derangement, cell roundness and neuro-vascular ingrowth.

**Fig. 1.**
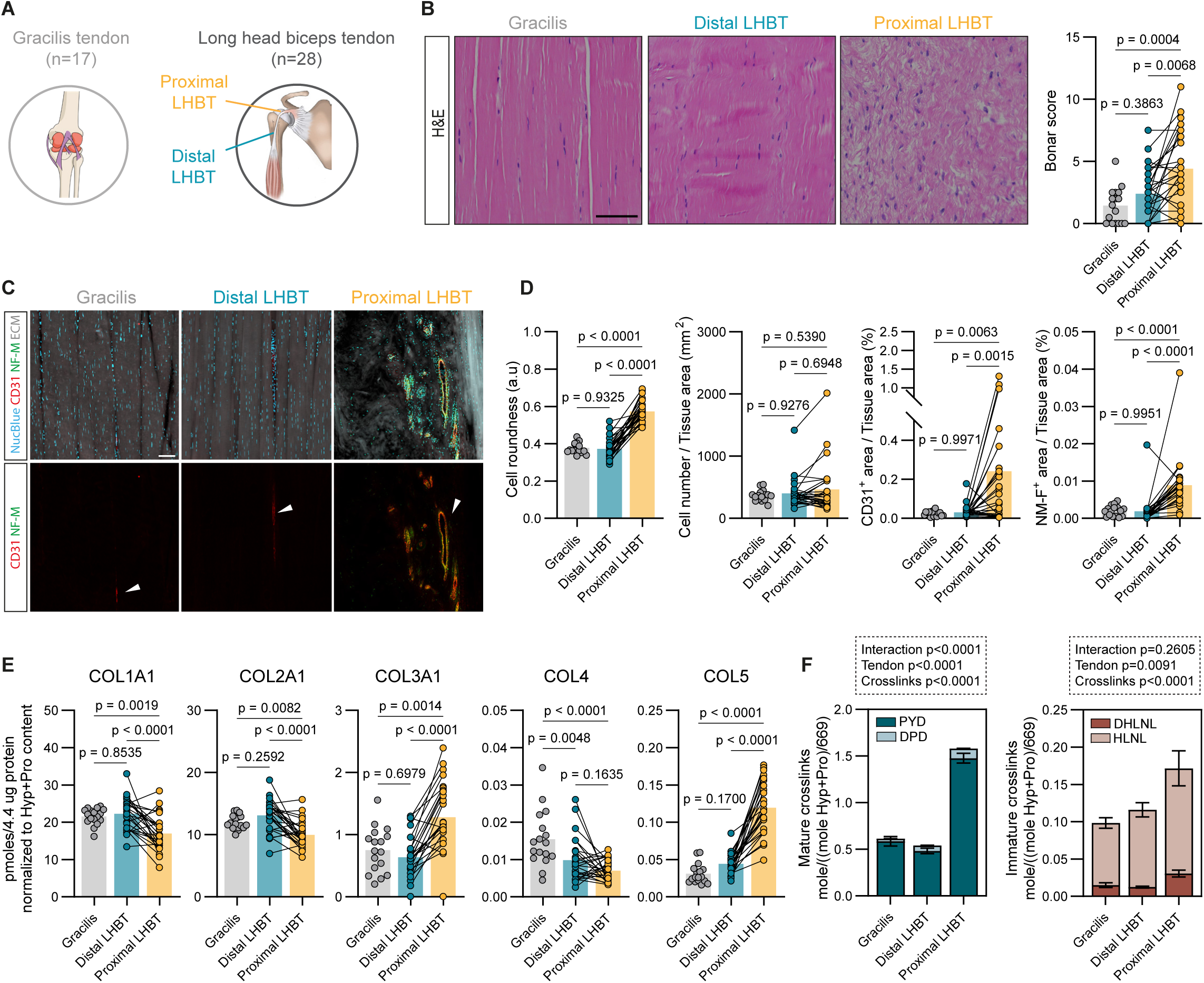
Diseased human tendons show pathological changes, characterized by ECM remodeling and increased collagen crosslinks. (**A**) Schematic representation of anatomical sourcing from two human cohorts. Healthy gracilis tendons were harvested as graft material from patients undergoing anterior cruciate ligament (ACL) or medial patellofemoral ligament (MPFL) reconstruction (n=17) while diseased long head biceps tendons were harvested from pathological shoulder joints (n=28). (**B**) Representative images of haematoxylin-eosin (H&E)-stained sections (left panel) and Bonar score (right panel) in human healthy gracilis (n=17) and diseased long head biceps tendons (n=28). Scale bar: 50µm. (**C**) Representative images of human healthy gracilis (n=17) and diseased long head biceps tendons (n=28) stained for the endothelial cells marker (CD31, red), myelinated nerve cells (NM-F, green), cell nuclei (NucBlue, cyan). Autofluorescence signal detected in the 488 channel was used to visualize ECM (grey). Scale bar: 100 µm. (**D**) Quantification of cell roundness and number (first and second plot, respectively), blood vessel (third plot) and myelinated nerve cell ingrowth (fourth plot) in the tendon core from panel C. (**E**) Mass-spectrometry based peptide quantification of COL1A1, COL2A1, COL3A1, COL4 and COL5 in gracilis (n=17) and diseased long head biceps tendons (n=26). Collagen peptides are expressed in picomoles per 4.4 microgram total protein normalized to the sum of hydroxyproline and proline (Hyp+Pro) content measured by amino acid analysis. (**F**) Mature and immature crosslinks levels normalized by the sample Hyp+Pro content and divided by total number of Hyp+Pro residues in collagen I (699) in human healthy gracilis (n=17) and diseased long head biceps tendons (n=26). Data are shown as mean with individual points representing biologically independent samples and lines connecting paired samples, except for panel F where data are mean ± SEM. One-way ANOVA with Tukey’s multiple comparisons test was used for statistical analysis in B, D and E. Two-way ANOVA (tendon, crosslinks) with Tukey’s multiple comparisons test was used in F. Created with Biorender.com.

## Hypoxia signaling is activated in human tendinopathy

Next, we investigated the biological processes underlying the observed pathological phenotype using an unbiased transcriptomics approach. Hence, we performed bulk RNA-seq of diseased proximal LHB samples and control distal LHB samples (n=15). Principal component analysis (PCA) revealed a clear separation between the two groups (**Fig. 2A**). A total of 1492 differentially expressed genes (DEGs; adjusted p value < 0.05 & abs (log2FC)> 1) were found, out of which 641 were upregulated and 851 were downregulated in the proximal LHB group compared to the control group (**Fig. 2B**). To characterize the upregulated DEGs in the diseased condition (Cluster 1), we performed pathway analysis using Hallmark (^40^), Reactome (^41,42^) and TRRUST (Transcriptional Regulatory Relationships Unraveled by Sentence-based Text mining) (^43^) databases (**Fig. 2C**). The results revealed a positive enrichment of epithelial mesenchymal transition (EMT), glycolysis, hypoxia and angiogenesis hallmark pathways. Reactome dataset analysis showed enrichment of collagen biosynthesis/assembly-related pathways and ECM organization, suggesting matrix turnover (**Fig. 2D and fig. S2A-D**). Additionally, TRRUST analysis identified enrichment of the HIF1α transcriptional factor (TF), along with other hypoxic- and EMT-related genes, such as VHL, ARNT and SMAD1 (**Fig. 2E**). The prominent enrichment of glycolysis, hypoxia and HIF1α in the diseased group prompted us to corroborate whether the HIF1α transcriptional response was activated in human tendon disease. Thus, we screened the canonical HIF1α target genes from Semenza gene set (^44^) and found that many of them, including *CA9, VEGFA* and *BNIP3,* were upregulated (adjusted p value <0.05) in the diseased group compared to the control (**Fig. 2F**). Immunofluorescence analysis confirmed increased HIF1α at the protein level, with strong nuclear HIF1α localization in the proximal LHB samples compared to LHB distal and gracilis samples, where HIF1α signals were weaker and primarily located in the cytoplasm (**Fig. 2G**). Furthermore, we verified these findings with independent samples. We performed label-free proteomics of diseased human rotator cuff tear tendons (RCTT, n=4) and healthy gracilis tendons (n=4). This analysis confirmed a positive enrichment of hypoxia and glycolysis signatures in the diseased group compared to the healthy controls, suggesting that this transcriptional factor may broadly underlie tendon pathology (**Fig. 2H and fig. S2E-G**).

**Fig. 2.**
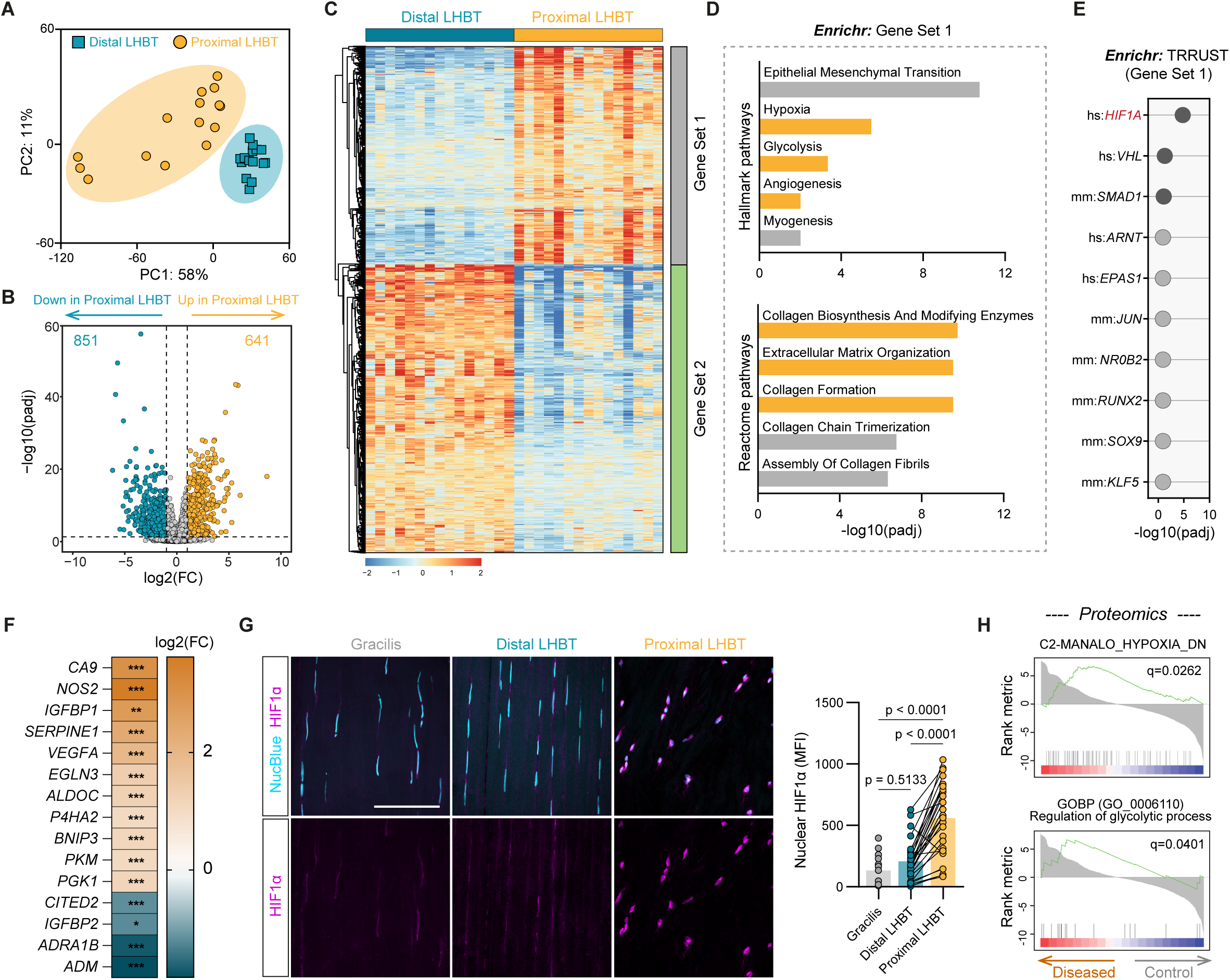
Transcriptomics analysis reveals ECM turnover and HIF1α upregulation in diseased human tendons. (**A**) Principal component analysis showing separation of proximal LHBT samples from the intra-patient distal LHBT samples (n=15). (**B**) Volcano plot of differentially expressed genes (DEGs) in diseased LHBT proximal region relative to the distal LHBT control. Colored dots show the 1492 significantly expressed genes, with the horizontal line corresponding to a padj<0.05 and vertical lines are at a cutoff of log2FC±1. (**C**) Heatmap of centered normalized counts of the whole 1492 DEGs in proximal LHBT samples versus distal LHBT controls. Columns represent individual samples (n=15). Gene Set 1 encloses the 641 upregulated DEGs; Gene Set 2 encloses the 851 downregulated DEGs. (**D**) Enriched pathway analysis of Gene Set 1 using *Enrichr* queried against MSigDB Hallmark and Reactome databases. Yellow bars highlight the hypoxic- and ECM-related pathways of interest upregulated in proximal LHBT samples. (**E**) Top 10 TF and non-TF target genes regulating Cluster 1 predicted by TRRUST database. The darker dots represent the significant TF and non-TF target genes; padj<0.05 (**F**) HIF1α downstream target genes from Semenza database which are significantly upregulated and downregulated (padj<0.05 and abs(log2(FC)>1) in the proximal LHBT group compared to the distal LHBT control. (**G**) Representative images (left panel) and quantification (right panel) of human gracilis (n=17) and long head biceps tendons (n=28) stained for HIF1α (magenta) and cell nuclei (NucBlue, cyan). Scale bar: 50 µm. Bar plots indicate the mean with individual points representing biologically independent samples and lines connecting paired samples. One-way ANOVA with Tukey’s multiple comparisons test was used for statistical analysis. (**H**) Gene set enrichment analysis (GSEA) of hypoxia and regulation of glycolytic process pathways conducted on label-free proteomics data of rotator cuff tear tendons (Diseased group) versus the gracilis control (Control group).

## Activation of HIF1α in tendon cells is sufficient to drive tendon disease in mice

Based on our observations of human tissues, we aimed to investigate a potentially causal role of HIF1α activation in tendon pathology. To this end, we generated mice with a tendon cell-targeted knockout of von Hippel-Lindau (VHL) protein by intercrossing *Scx*^Cre^ mice, in which Cre recombinase is under the *Scleraxis* promoter, with mice carrying the loxP -flanked *Vhl* alleles (**Fig. 3A**). VHL acts as a ubiquitin ligase that marks HIFs for proteolytic degradation under normoxic conditions, thereby silencing the HIF-dependent transcriptional program. Hence, *Vhl* knockout results in stabilization of HIFs (^45^). We found increased HIF1α and HIF2α protein levels in the core tenocytes of *Scx*^Cre^;*Vhl*^f/f^ mice (**Fig. 3B and fig. S3A-C**). This resulted in the activation of HIF-dependent angiogenic and metabolic transcriptional programs (**fig. S3D-J**). To assess how HIFs activation affects tendon structure and function, we conducted a comprehensive characterization of *Scx*^Cre^;*Vhl*^f/f^ tendons. Histological analysis of *Scx*^Cre^;*Vhl*^f/f^ Achilles tendons showed pronounced similarities to human tendinopathy (**Fig. 1D)**, including increased cell number and roundness (**Fig. 3C**). Confocal images of *Scx*^Cre^;*Vhl*^f/f^ mice crossed with *Scx*GFP reporter line, demonstrated that the shift from spindle to round shape occurred in tenocytes (*Scx*GFP+ cells) (**Fig. 3D and fig. S7A**). These pathological changes were accompanied by collagen fiber disorganization (**Fig. 3E**) and altered ECM composition (**Fig. 3F**), consistent with the matrix dysregulation observed in human samples. Indeed, proteome profile of *Scx*^Cre^;*Vhl*^f/f^ tail tendons revealed increased levels of COL3A1, COL5A1 and COL5A2, and decreased levels of COL1A1 and COL1A2. Bulk RNA-seq further confirmed these changes, showing upregulation of *Col3a1/Col1a1* ratio and increase expression of matrix metalloproteinases (*Mmp9*, *Mmp3*, *Mmp13*) in the *Vhl-KO* mice tail tendons, indicating elevated matrix turnover (**Fig. 3G and fig. S4A**). Because impaired collagen synthesis in the human cohort was associated with a fibrotic phenotype (**Fig. 1E and F**), we again looked at the crosslink content and found that *Scx*^Cre^;*Vhl*^f/f^ tail tendons exhibited a similar increase in mature crosslinks as well as in immature PYD- precursors DHLNL crosslinks, whereas the immature DPD-precursors HLNL amount was almost halved compared to the WT littermates (**Fig. 3H and fig. S4B-C**). An increase was also observed in lysyl oxidase (LOX) activity and lysyl hydroxylase 2 (PLOD2) levels of *Scx*^Cre^;*Vhl*^f/f^ Achilles and tails tendons, two key enzymes involved in collagen crosslink formation (^46^) (**fig. S4D and E**). To assess whether this increased collagen crosslinking resulted in a stronger and stiffer tendon (^47,48^), we performed mechanical testing. Interestingly, despite more extensive crosslinking, *Scx*^Cre^;*Vhl*^f/f^ tendon fascicles showed a 33% reduction in the elastic (E) modulus and an increased failure stress (**Fig. 3I and fig. S4F**). In addition, stress-relaxation experiments revealed decreased viscoelasticity, with a 30% decrease of stress decay (**Fig. 3J and fig. S4G**), characterizing tissue fibrosis (^49,50^) which sufficed to increase the calcium activation threshold (**fig. S4H**). We then asked whether *Vhl*-deficiency in tendon cells could lead to the neovascularization and innervation of the tendon proper that characterizes chronic human tendinopathy. (**Fig. 1C and D**). High-resolution 3D scans of whole-mount *Vhl*-deficient mouse Achilles tendons showed a dense vascular network in the tendon core (**Fig. 3K**) associated with higher presence of myelinated nerve fibers (**Fig. 3L**). Additionally, we confirmed an increased CD31^+^ cell population in tail tendons by FACs analysis (**fig. S5**).

**Fig. 3.**
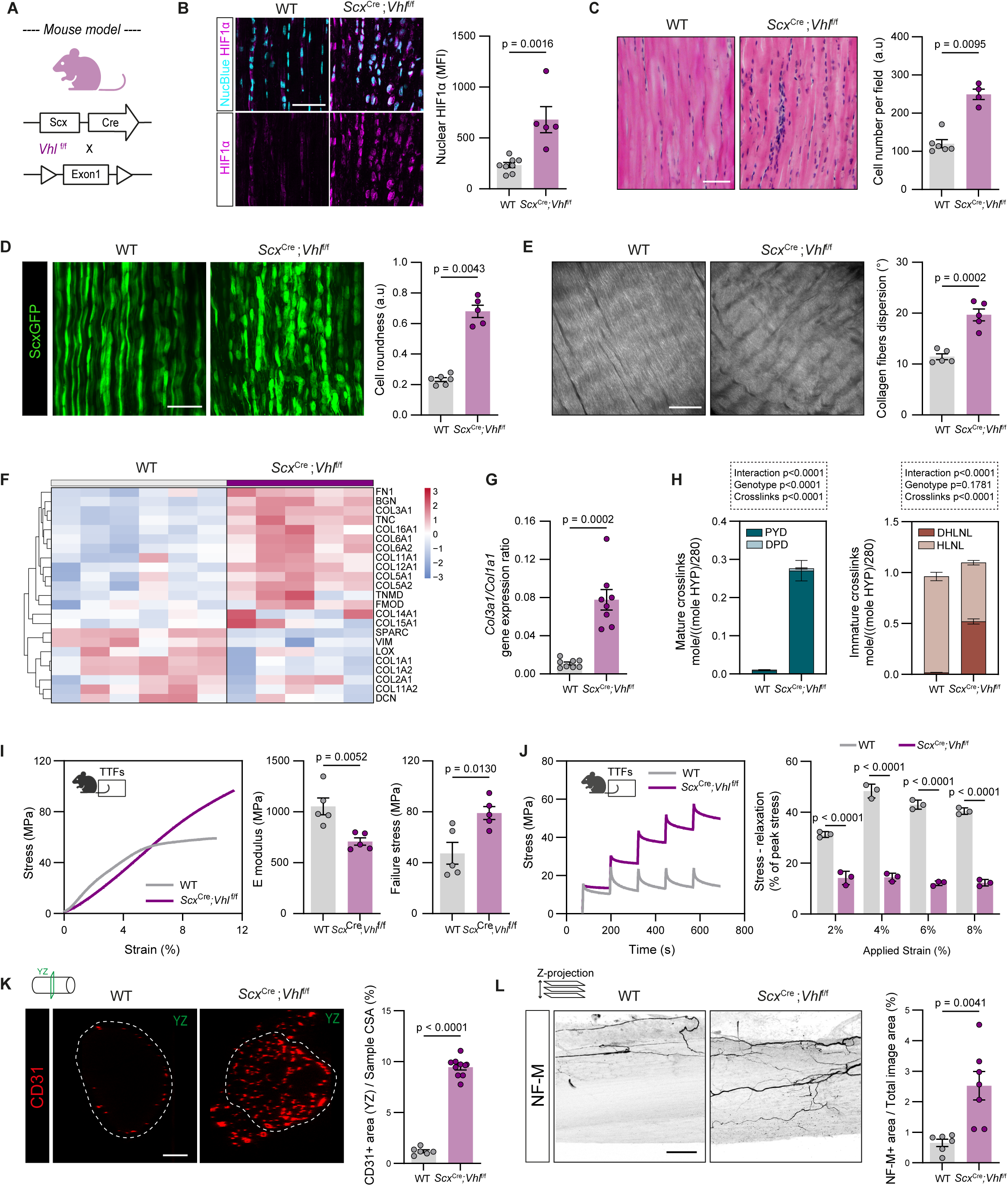
HIFs activation in murine tendon fibroblasts results in tendon pathological features. (**A**) Generation of *Scx*^Cre^;*Vhl*^f/f^ mice expressing constitutive Cre under the *Scx* promoter. (**B**) Representative images (left panel) and quantification (right panel) of HIF1α nuclear translocation based on colocalization with NucBlue staining in WT (n= 8) and *Scx*^Cre^;*Vhl*^f/f^ (n=5) Achilles tendon. Scale bar: 50µm. (**C**) Representative images of haematoxylin-eosin (H&E)-stained sections (right panel) and cell number quantification (left panel) in WT (n=6) and *Scx*^Cre^;*Vhl*^f/f^ (n=4) Achilles tendon. Scale bar: 100µm. (**D**) Representative images (left panel) and cell roundness quantification (right panel) of whole mount *Scx*^GFP+^ cells in WT (n=6) and *Scx*^Cre^;*Vhl*^f/f^ (n=5) Achilles tendon. Scale bar: 50µm. (**E**) Representative images of second harmonic generation images (left panel) and collagen dispersion quantification of WT (n=5) and *Scx*^Cre^;*Vhl*^f/f^ (n=5) Achilles tendon. Scale bar: 50µm. (**F**) Heatmap of ECM-related proteins expressed as centered normalized intensities in WT (n=6) and *Scx*^Cre^;*Vhl*^f/f^ (n=5) TTFs. (**G**) *Col3a1*/ *Col1a1* ratio calculated from bulk RNA-seq normalized counts in WT (n=8) and *Scx*^Cre^;*Vhl*^f/f^ (n=8) tail tendon fascicles. (**H**) Mature and immature crosslinks levels normalized by the sample Hyp content divided by total number of Hyp residues in collagen I (280) in WT (n=4) and *Scx*^Cre^;*Vhl*^f/f^ (n=4) TTFs. Two-way ANOVA with Sidak’s multiple comparisons test was used for statistical analysis (**I**) Ramp-to-failure test shows a decreased E modulus and increased failure stress in *Scx*^Cre^;*Vhl*^f/f^ (n=5) TTFs compared to the WT littermates (n=5). n=6 fascicles tested per mouse. (**J**) Stress-relaxation test reveals a reduced stress decay in *Scx*^Cre^;*Vhl*^f/f^ (n=3) TTFs compared to the WT littermates (n=3). n=4 fascicles tested per mouse. (**K**) Representative images (left panel) and quantification (right panel) of ECs (CD31, red) in the YZ-plane of whole-mount Achilles tendon in *Scx*^Cre^;*Vhl*^f/f^ (n=10) compared to the WT controls (n= 6). CD31^+^ area is normalized by sample cross-sectional. Scale bar: 50µm. (**L**) Representative images (left panel) and quantification (right panel) of myelinated nerve cells (NM-F, black) in *Scx*^Cre^;*Vhl*^f/f^ (n=7) and WT controls (n= 6) whole-mount Achilles tendon visualized and quantified as maximum intensity projection. NM-F ^+^ area is normalized by total image area. Scale bar: 50µm. Each point represents a single mouse. Bar graphs indicate mean ± SEM. Student’s t test (unpaired, two tailed) was used in B,C,D,E,G,I,K,L. Two-way ANOVA with Tukey’s multiple comparisons test was used in H,J.

Given that *Vhl* deletion results in HIF1α and HIF2α stabilization, we next asked whether the observed phenotype is specifically HIF1α dependent. Thus, we co-silenced *Vhl* and *Hif1α* by intercrossing the *Scx*^Cre^;*Vhl*^f/f^ mice with *Hif1a*^f/f^. Indeed, *Scx*^Cre^;*Vhl*^f/f^;*Hif1α*^f/f^ mice showed normalization of all pathological features except for hypercellularity, which was not rescued and thus might be attributed to HIF2α stabilization (**fig. S6**). Collectively, these data support the notion that HIF1α activation in the core tenocytes suffices to drive key hallmarks of tendon pathology.

## VEGFA-mediated vascular recruitment drives tissue innervation but is not required to provoke pathological tendon histology, matrisome or mechanical properties

Hypervascularity is a central hallmark of tendinopathy and has been widely discussed as a driver of tendon disease and a potentially powerful point for clinical intervention (^21^). We sought to test this hypothesis in our mouse model by decoupling the effect of HIF1α activation from blood vessel ingrowth using our *Scx*^Cre^;*Vhl*^f/f^ mouse model. We first FACS sorted *Scx*^GFP+^ tendon cells from WT and *Scx*^Cre^;*Scx*GFP;*Vhl*^f/f^ tail tendon fascicles (TTFs) and screened for potential angiogenic cytokines involved in EC recruitment. Immunoassay analysis of conditioned media from *GFP+-Vhl-KO* tenocytes revealed numerous overexpressed pro-angiogenic factors compared to the WT, among which VEGFA was the most upregulated one with over 4-fold increase in concentration (**Fig. 4A and fig. S7**). Furthermore, ECs spheroids treated with human *VHL-KO* tendon cell-derived conditioned media showed increased EC sprouting compared to controls (**fig. S8**). We thus generated a double knockout mouse model by intercrossing the *Scx*^Cre^;*Vhl*^f/f^ mice with *Vegfa*^f/f^ mice to investigate the potentially causal contribution of VEGFA to VHL-driven tendinopathy (**Fig. 4B**). Deletion of one *Vegfa* allele was sufficient to prevent ingrowth of blood vessels in *Scx*^Cre^;*Vhl*^f/f^;*Vegfa*+/f Achilles tendon, confirming that vascular recruitment is VEGFA-driven. Interestingly, we also observed that deleting VEGFA reduced the presence of myelinated nerves, suggesting a neurovascular link in tendinopathy (**Fig. 4C and D**). Histological analysis of the double-KO Achilles tendon showed no overt differences in cell roundness compared to the *Vhl-KO* samples, although a trend towards a lower cell number was observed (**Fig. 4E**). In contrast, label-free proteomics showed that loss of *Vegfa* failed to restore the ECM-related proteins which were differentially expressed in the *Vhl-KO* tendons, such as COL3A1, COL5A1, COL5A2, TNMD and BGN even though we observed an attenuation (**Fig. 4F**). Furthermore, ramp-to-failure and stress-relaxation tests did not show a restoration of E modulus, failure stress and viscoelasticity in the *Scx*^Cre^;*Vhl*^f/f^;*Vegfa*+/f tendons (**Fig. 4G and H**). In addition, *Scx*^Cre^;*Vhl*^f/f^;*Vegfa*+/f tendons showed comparable amounts of LOX activity and crosslinks to the *Scx*^Cre^;*Vhl*^f/f^ mice, except for the total glycation which was reduced to the WT levels (**Fig. 4I and fig. S9**). Taken together, these findings reveal that HIF1α stabilization and activation promotes angiogenesis and innervation in the tendon core via VEGFA. However, in our mouse model, blood vessel ingrowth is not responsible for the observed phenotypic features of tendinopathy, including cell roundness, matrisome heterogeneity and altered mechanics.

**Fig. 4.**
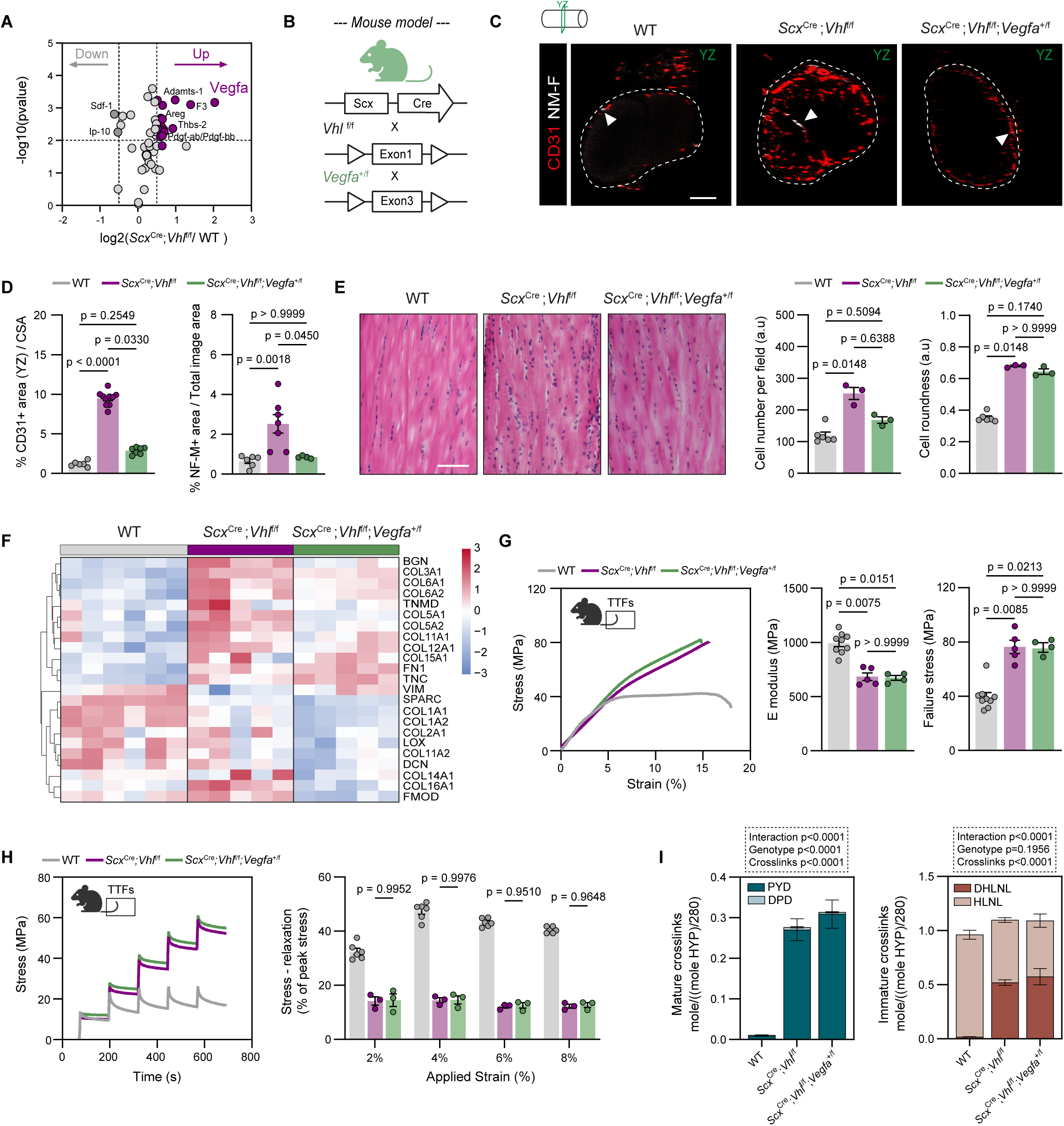
HIF1α-driven alterations in tendon histology, matrisome and mechanical properties do not depend on vascular recruitment via VEGFA secretion. (**A**) Volcano plot of significant upregulated (violet) and downregulated (grey) pro-angiogenic cytokines in tenocytes conditioned media of *Scx*^Cre^;*Vhl*^f/f^ TTFs compared to WT control. Horizontal line corresponding to a pvalue<0.01 and vertical lines are at a cutoff of log2(*Scx*^Cre^;*Vhl*^f/f^/WT)±0.5. (**B**) Generation of *Scx*^Cre^;*Vhl*^fl/fl^;*Vegfa*^+/fl^ double knockout mice expressing constitutive Cre under the *Scx* promoter. (**C**) Representative whole-mount images of wild type, *Scx*^Cre^;*Vhl*^f/f^ and *Scx*^Cre^;*Vhl*^fl/fl^;*Vegfa*^+/fl^ Achilles tendon stained for ECs (CD31, red) and myelinated nerve cells (NM-F, white) markers. Images show tendon cross-sectional area (YZ plane). Whyte arrows point at the myelinated nerve cells. Scale bar: 100µm. (**D**) Quantification of ECs (CD31^+^) in WT controls (n= 6), *Scx*^Cre^;*Vhl*^f/f^ (n=10) and *Scx*^Cre^;*Vhl*^fl/fl^;*Vegfa*^+/fl^ (n=7) relative to the tendon CSA and myelinated nerve cells (NM-F+) in WT controls (n= 6), *Scx*^Cre^;*Vhl*^f/f^ (n=7) and *Scx*^Cre^;*Vhl*^fl/fl^;*Vegfa*^+/fl^ (n=5). NM-F^+^ signal was quantified as maximum intensity projection normalized to total image area. (**E**) Representative images of haematoxylin-eosin (H&E)-stained sections (right panel) in WT (n=6), *Scx*^Cre^;*Vhl*^f/f^ (n=3) and *Scx*^Cre^;*Vhl*^fl/fl^;*Vegfa*^+/fl^ (n=3) Achilles tendon. Quantification of cell number per field of view and cell roundness (left panel). Scale bar: 50µm. (**F**) Heatmap of ECM-related proteins expressed as centered normalized intensities in WT (n=6), *Scx*^Cre^;*Vhl*^f/f^ (n=5) and *Scx*^Cre^;*Vhl*^fl/fl^;*Vegfa*^+/fl^ (n=5). (**G**) Ramp- to-failure test shows a decreased E modulus and increased failure stress in both *Scx*^Cre^;*Vhl*^f/f^ (n=5) and *Scx*^Cre^;*Vhl*^fl/fl^;*Vegfa*^+/fl^ (n=4) TTFs compared to the WT littermates (n= 9). n=6 fascicles tested per mouse. (**H**) Stress-relaxation test reveals a similar reduction of stress decay in *Scx*^Cre^;*Vhl*^f/f^ (n=3) and *Scx*^Cre^;*Vhl*^fl/fl^;*Vegfa*^+/fl^ (n=3) TTFs compared to the WT littermates (n= 6). n=4 fascicles tested per mouse. (**I**) Mature and immature crosslinks levels normalized by the sample Hyp content divided by total number of Hyp residues in collagen I (280) in WT (n=10), *Scx*^Cre^;*Vhl*^f/f^ (n=4) and *Scx*^Cre^;*Vhl*^fl/fl^;*Vegfa*^+/fl^ (n=6). Each point represents a single mouse. Bar graphs indicate mean ± SEM. Kruskal-Wallis with Dunn’s multiple comparison test was used in D,E,G. Two-way ANOVA with Tukey’s multiple comparisons test was used in H,I.

## Mechanical overload induces HIF1α activation

Next, we sought to investigate whether HIF1α activation is under mechanical control in tendon cells. To this end, we customized a bioreactor system to enable cyclic, uniaxial tensile stretching of human-derived tenocytes seeded on a silicone chamber (**Fig. 5A and fig. S10A-C**). Cellular strain levels associated with overload were based on mechanically activated calcium (Ca^2+^) signaling which we previously identified as a physiological ‘limit switch’ related to the onset of tendon damage upon overload (^51^) (**Fig. 5B and fig. S10D**). To assess the time-dependent response of HIF1α to mechanical load, we stimulated cells with strain parameters below (low load) and above (overload) the Ca^2+^ signaling threshold, and measured HIF1α protein levels after 1h, 2h, 4h and 6h of continuous stretching (**fig. 5A**). As a control, cells were placed in the silicone chamber with no stimulation. Mild changes in HIF1α levels were detected after 1h and 2h of stimulation in the low load condition, with an observed 2- fold change increase in HIF1α signal at 4h, which diminished at 6h. In contrast, the overload stimulation triggered a robust HIF1α response as soon as after 1h of stimulus, with a steep increase at 2h, a peak at 4h and a moderate decreased at 6h (**Fig. 5C-D and fig. S10E**). Similar results were identified upon nuclear fractionation using an ELISA which detects HIF1α DNA-binding ability. This confirmed HIF1α nuclear translocation and activation upon mechanical stimulation in a strain-dependent manner (**Fig. 5E**). Furthermore, using the same experimental protocol, we found a significant increase in VEGFA secretion at 72h post-stretching (**Fig. 5F and G**). Taken together those findings suggest that mechanical overload leads to HIF1α stabilization and activation of its downstream target VEGFA.

**Fig. 5.**
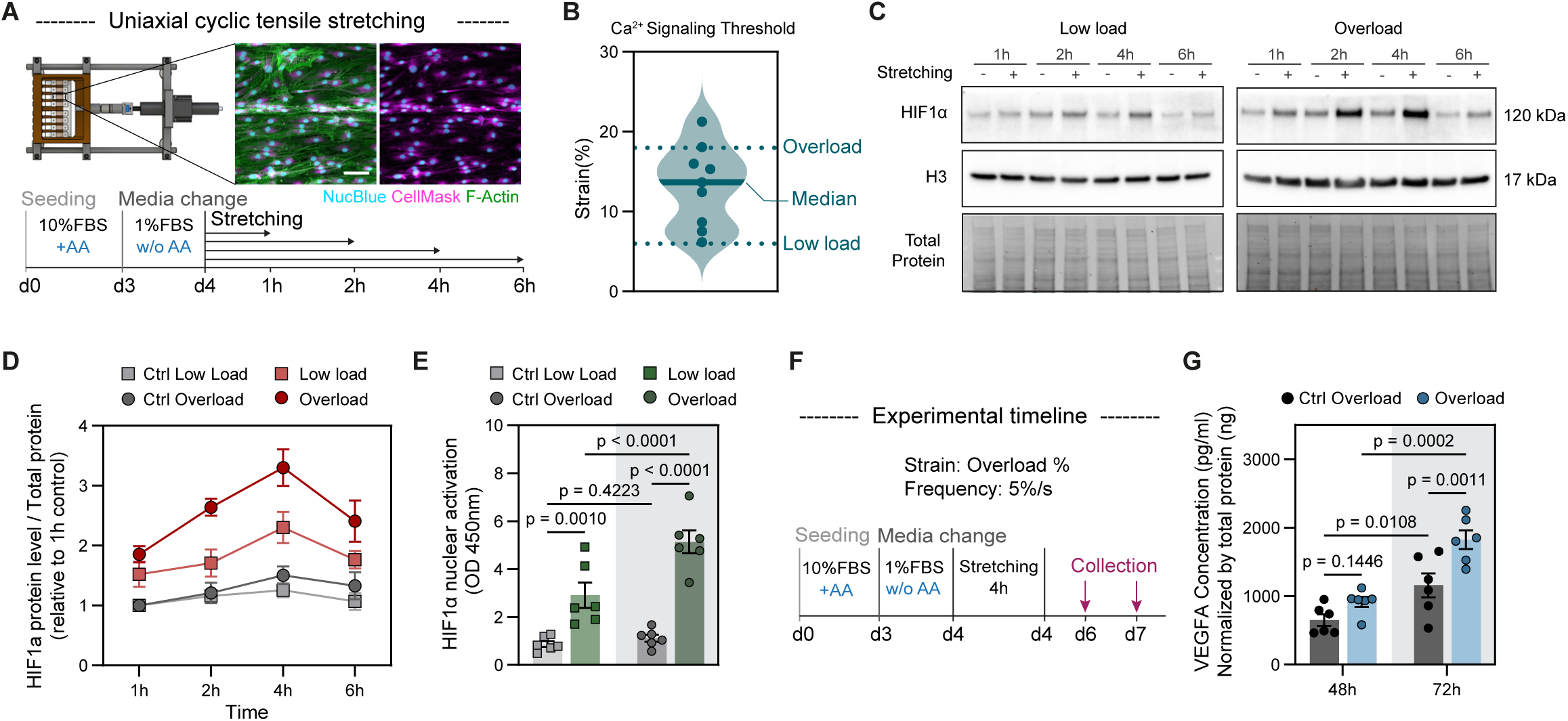
HIF1α activation in response to mechanical stretching. (**A**) Schematic representation of the technical set up and experimental timeline. Human-derived tendon fibroblasts seeded on silicone chambers are stained for F-actin (Phalloidin, green), plasma membrane (CellMask, magenta) and cell nuclei (NucBlue, cyan). Scale bar: 100µm. (**B**) Quantification of human tenocytes Ca^2+^ signaling threshold upon stretching after 3 days of culture in silicone chambers. Replicates are biological (n=9 donors), and they represent the average of three technical replicates (n=27 chambers). (**C**) Time-dependent HIF1α signal and (**D**) quantification of human tenocytes upon cyclic, uniaxial tensile stretching at two different strain percentages (n=6). Controls and stretched samples are normalized to 1h control. (**E**) ELISA quantification of nuclear HIF1α activation in human tenocytes subjected to stretched and overstretched mechanical stimulations at 4 hours (n=6). (**F**) Schematic representation of experimental timeline.(**G**) VEGFA concentration in human tendon cell-conditioned media, measured by ELISA at 48h and 72h post-stimulation, normalized to protein concentration (n=6). Data points represent biologically independent samples. Bar graphs indicate mean ± SEM. Two-way ANOVA with Tukey’s multiple comparisons test was used in D,E,G.

## HIF1α inhibition rescues overload- induced tendon maladaptation in mice

We next aimed to elucidate whether HIF1α activation may act as a checkpoint in response to overload *in vivo*. To this end, we employed a surgical model of synergist ablation (SynAb), where the Achilles tendon complexes are ablated, resulting in substantial overload of the plantaris tendons (^52^) (**Fig. 6A**). This model has recently been described to induce tendon degeneration within 8 weeks post-surgery (^53^). To assess the effects of extended overload beyond 8 weeks, we first characterized pathological changes in B6/J mice plantaris tendons at 2 and 12 weeks post-surgery (**fig. S11**). Those experiments confirmed significant structural and functional impairment in the plantaris tendon after 12, but not 2, weeks of prolonged overload (**fig. S11**). We thus decided to perform the same surgery in WT and *Scx*^Cre^;*Hif1α*^f/f^ mice, and studied tendon properties after 12 weeks of overload. Under baseline (sham) conditions, no overt differences in diameter, histology, ECM, mechanical properties and crosslinks were found between the WT and *Scx*^Cre^;*Hif1α*^f/f^ plantaris tendons. Overloaded plantaris tendon cores of both wild type and *Scx*^Cre^;*Hif1α*^f/f^ mice showed an 11% increase in diameter (**Fig. 6B**). But while histological analysis of the WT-loaded plantaris (SynAb WT) tendons confirmed the development of several tendinopathic hallmarks such as hypercellularity, increased cell roundness and an accumulation of cell ‘pockets’ between collagen fibers, removing HIF1α effectively rescued the phenotype, reducing the cell number and roundness to the sham levels (**Fig. 6C and D**). Qualitative assessment of collagen fibers size with picrosirius red staining (PSR) under polarized light revealed significant changes in the SynAb WT group, characterized by development of spectral signatures associated with smaller /immature collagen fibers (^54^) that were absent in the SynAb *Scx*^Cre^;*Hif1α*^f/f^ samples (**Fig. 6E**). As tendon mechanical properties reflect tissue structural and functional integrity, we next performed ramp-to-failure testing and found a 19% decrease in stiffness and a 22% increase in failure strain in the SynAb WT plantaris tendon, indicating altered mechanical properties. In contrast, we did not observe these changes in the SynAb *Scx*^Cre^;*Hif1α*^f/f^ samples, which maintained similar stiffness and failure strain relative to the sham control tendons (**Fig. 6F**). Finally, crosslink analysis revealed a 2-fold increase in PYD crosslinks in the SynAb WT at 12 weeks, while the knockout group did not exhibit this increase (**Fig. 6G).** The same trend was observed for the PYD-precursor DHLNL crosslinks (**fig. S12**). Collectively, these findings suggest that HIF1α inhibition in tendon cells prevents tendon maladaptation in response to overload.

**Fig. 6.**
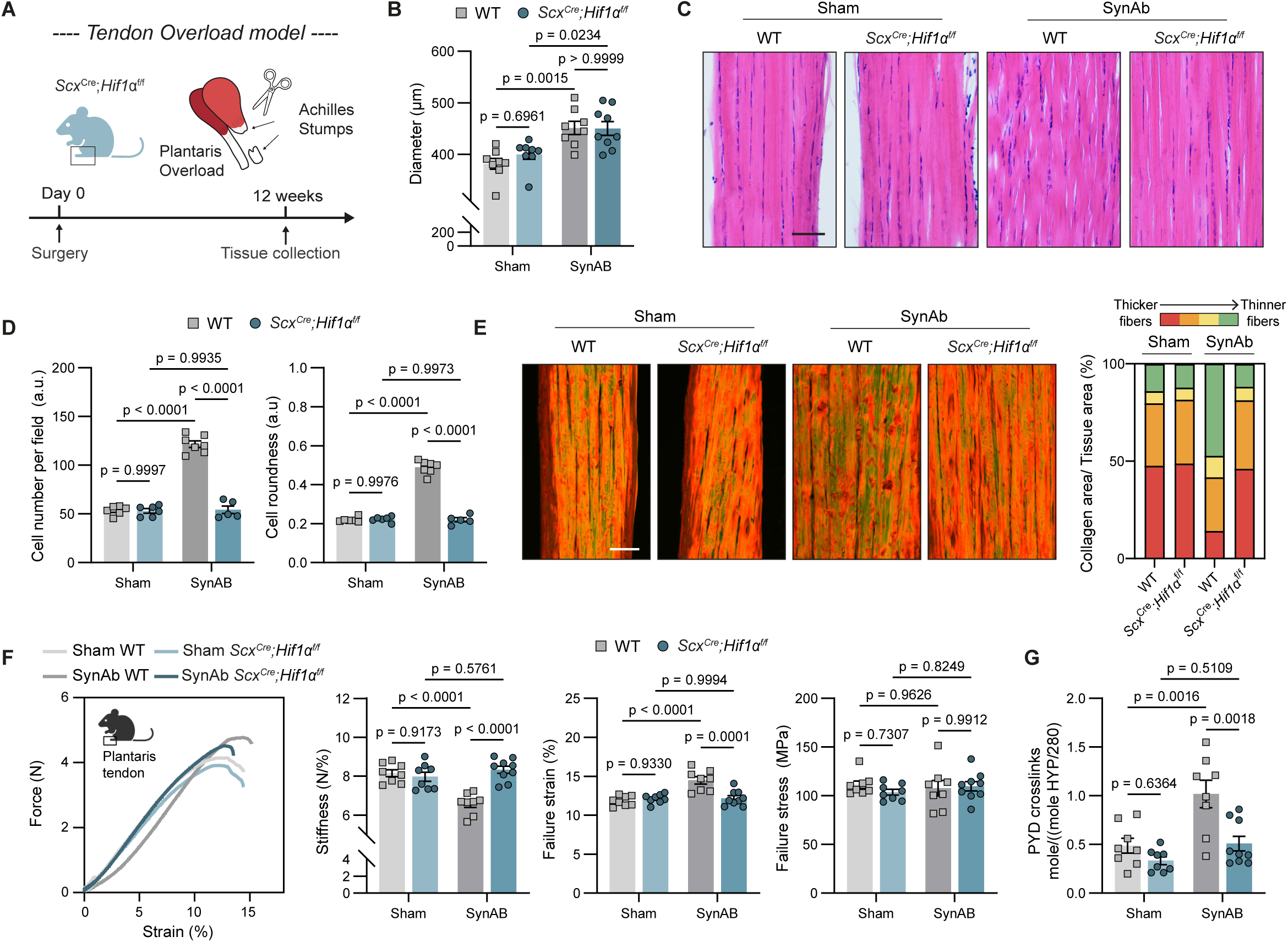
Knockout of HIF1α in tendon cells rescues tendon maladaptations in response to overload. (**A**) Experimental schematic of the synergist ablation (SynAb) model. (**B**) Quantification of plantaris tendon diameter at 12 weeks post-surgery of WT (n=8, Sham; n=8, SynAb) and *Scx*^Cre^;*Hif1α*^f/f^ (n=7, Sham; n=8, SynAb) mice. (**C**) Representative images of haematoxylin-eosin (H&E)-stained sections and (**D**) cell number and cell roundness quantification of WT (n=7, Sham; n=6, SynAb) and *Scx*^Cre^;*Hif1α*^f/f^ (n=7, Sham; n=5, SynAb) plantaris tendon. Scale bar: 50µm. (**E**) Representative images of picrosirius red (PSR) staining under polarized light and qualitative quantification of collagen fibers content in WT (n=6, Sham; n=6, SynAb) and *Scx*^Cre^;*Hif1α*^f/f^ (n=7, Sham; n=5, SynAb) plantaris tendon. Red, orange, yellow and green colors represent different types of collagen fibers based on their thickness. Red indicates thicker/mature collagen fibers while green indicates thinner/ immature fibers. Scale bar: 50µm. (**F**) Ramp-to-failure test shows a decreased stiffness and increased failure strain in the SynAb WT group which is normalized to the Sham levels in the SynAb *Scx*^Cre^;*Hif1α*^f/f^ group. n=9,Sham; n=9,SynAb for the WT conditions and n=8,Sham; n=9,SynAb for the *Scx*^Cre^;*Hif1α*^f/f^ groups. (**G**) Mature PYD crosslinks levels normalized by the sample Hyp content divided by total number of Hyp residues in collagen I (280) in the WT (n=8, Sham; n=8, SynAb) and *Scx*^Cre^;*Hif1α*^f/f^ groups (n=8 Sham; n=9 SynAb). Data points represent biologically independent samples. Bar graphs indicate mean ± SEM. Two-way ANOVA with Tukey’s multiple comparisons test was used in B,D, F,G, H,I

## DISCUSSION

Chronic tendinopathy is a disease condition characterized by extracellular matrix derangement, hypercellularity, fibrotic hallmarks, and neuro-vascular ingrowth. How this pathological switch is initiated remains unclear, as does the role of vascularization in tendon pathology. Our study identifies HIF1α, a transcriptional regulator of the cellular response to hypoxia (^55^), as a pivotal checkpoint for tendon maladaptation in response to mechanical overload. We show that HIF1α is activated after mechanical overload and, when chronically activated, triggers pathological hallmarks of tendinopathy independently of vascular recruitment. Notably, inhibition of HIF1α rescues these adverse effects, suggesting its potential as a therapeutic target.

Previous research has associated HIF1α with several degenerative processes in tendinopathy, including increased apoptosis, inflammatory response, and collagen degradation in human tissue samples (^25,27,28,56–59^). The increased HIF1α levels reported in these studies was based on assessments of diseased rotator cuff tendons analyzed by immunohistochemistry and /or immunofluorescence staining. In our investigation, we confirmed enrichment of HIF1α signaling in diseased rotator cuff tendons by using transcriptomics, immunofluorescence, and label-free proteomics, further identifying elevated expression in diseased long head biceps tendon (LHBT) samples. Extensive characterization revealed that the proximal LHBT region displayed advanced pathological hallmarks associated with increased HIF1α levels, in contrast to the distal LHBT region and healthy gracilis samples from the same patients. These findings suggest that hypoxic signaling is a shared feature of both early- and late-stage tendinopathies and could be broadly relevant as a tendon pathomechanism.

Traditionally, the field has considered HIF1α expression as a cellular protective response within nutrient-poor tissue, gradually overwhelmed by overuse related damage until it descends into chronic pathology. However, our data unequivocally show that chronic HIF1α activation alone (following genetic deletion of *Vhl*) recapitulates key biophysical hallmarks of human tendinopathy, including aberrant collagen matrix structure and neuro-vascular recruitment. Mechanical function was severely compromised, with PYD collagen crosslinking levels more than three times higher than those in 100-week-old mice (^60^), accompanied by neuro-vascular invasion. These insights show that hypoxia signaling in tendon disease may not just be a downstream protective mechanism but is sufficient to drive disease onset and progression.

Furthermore, by decoupling vascular recruitment from HIF1α activation with a novel *Scx*^Cre^;*Vhl*^fl/fl^;*Vegfa*^+/fl^ mouse model, we revealed that aberrant matrix alterations due to HIF1α activation were independent of vascular changes. This finding answers a longstanding question in the field whether vascular invasion is requisite for the progression of tendinopathy (^16,61,62^). On one hand, these findings strongly challenge the rationale for targeting tendon vasculature therapeutically to resolve disease. On the other hand, future research needs to unravel whether HIF1α-driven neurovascular infiltration could play a key role in the clinical manifestation of pain in tendinopathy.

While our study clearly links HIF1α activation to tendon tissue derangement, the downstream chain of events behind this degenerative process remains somewhat unclear. Given the exquisite sensitivity of stromal cells to the viscoelasticity of their local matrix(^63–67^), we speculate that HIF1α-driven collagen crosslinking could be the beginning of this chain. This likely would include matrix driven transcriptional rewiring, remodeling of the pericellular matrix, cellular rounding and distancing from the collagen backbone, and proliferation. Indeed, our histological sections and multiomic datasets suggest that all of these processes are at work, with crosslink-induced loss of tissue viscoelasticity markedly affecting mechanotransduction (**fig. S4H**).

This loss of viscoelasticity could explain or coexist with mechanisms identified in prior studies that link HIF1α activation to collagen biosynthesis and crosslinking in chondrocytes via altered cellular metabolism (^68^). Our data consistently show a HIF1α- gated matrisome switch away from collagen type-I, the primary functional matrix constituent of tendon, in favor of other fibrillar collagens. Within this shift, the secretion of collagen type-III may be particularly relevant, as it triggers inflammatory pathways that can drive tendinopathy. Untangling how HIF1α-activated collagen crosslinking, matrix viscoelasticity, cellular mechanotransduction, cell-matrix interactions, and tissue crosstalk cooperate to drive the progression of tissue derangement represents a rich area for future research.

Upstream of HIF1α activation, our data directly link mechanical overload to HIF1α activation and HIF1α gated matrix maladaptation. In vitro, we mechanically loaded matrix-embedded, human-derived tendon cells beyond tissue strain limits known to robustly trigger stretch-activated calcium channels. This ’overloading’ resulted in a clear upregulation of HIF1α compared to constructs loaded below this threshold. This experiment advances beyond previous work of others (^51^) by explicitly tying mechanical stimulation to a contextual mechanotransduction event (calcium influx in response to matrix overstretch); however, it is nonetheless consistent with prior studies that have reported HIF1α stabilization following mechanical loading in different cell types (^69–72^) including *in vitro* mechanical loading studies on fibroblasts that have also directly linked mechanical loading to angiogenic and pro-inflammatory cytokines (^73–75^).

*In vivo*, we utilized a tendon overload model, employing the synergist ablation technique recently shown to induce tendon functional degeneration in response to chronic overload (^53^). Upon characterizing the overload-induced maladaptation at two distinct time points (**fig. S10**), we observed that the severity of tissue degeneration increased with the duration of sustained overload. After 2 weeks of synergist ablation, we detected thicker tendons with still well-organized collagen. We speculate that, at early time points, the overload-stimulated tenocytes become activated, initiating tissue remodeling and adaptation to sustain higher loads. By 12 weeks, however, mechanical and histological analyses revealed hallmarks of chronic tendinopathy, including collagen matrix derangement despite continued hypertrophy. Histopathological features, such as increased cell density, cellular rounding, and smaller collagen fibers, were accompanied by reduced mechanical stiffness. Notably, inhibition of HIF1α in tendon fibroblasts effectively prevented these pathological features. Our findings suggest that HIF1α downstream targets or upstream activating mechanisms could offer novel treatment strategies for clinical tendinopathy, potentially using antibody-displaying extracellular vesicles (^76^).

Our *in vivo* overload experiments also reveal important connections between mechanical overload, HIF1α and mechanisms of functional adaptation. In the 12 weeks following synergist ablation the overloaded plantaris tendons showed a two-fold increase in PYD crosslinks content compared to the sham group, with evident biomechanical alterations in both the elastic and viscoelastic behavior of the tendon.

In contrast, overloaded tendon from *Scx*^Cre^;*Hif1α*^f/f^ mice displayed PYD levels comparable to those of the sham WT mice. Despite a similar adaptive increase in cross sectional area in both groups, only the wild type mice with elevated crosslinking levels demonstrated a measurable adaptation of tendon biomechanical properties (**Fig. 6G**). In particular, tendon stiffness decreased in WT mice, despite the increased tendon cross-section, reflecting the diminished matrix quality observed in histological sections. Notably, the viscoelastic and failure behaviors of these tendons were measurably altered, in contrast to HIFα knockout tendons whose functional properties did not appreciably change despite clear tissue hypertrophy in response to overload.

Although our findings of chronic HIF1α activation in diseased human tendons confirm earlier studies, this is the first to provide a quantitative analysis of key collagen matrix species and crosslinking profiles in human tendinopathy. This novel dataset moves beyond the qualitative assessments typically employed in tendinopathy research, potentially offering a functionally relevant and sensitive means of monitoring changes in matrix composition. By quantifying specific collagen isoforms and their crosslinking patterns, our data provide critical insights into tendon matrix biomechanical status and remodeling history, particularly in relation to aberrant matrix turnover. Unlike previous studies, which have largely focused on immunohistochemical and morphological evaluations, our use of label-free proteomics allows for a more nuanced interpretation of how changes in matrix composition directly impact tissue function. The ability to track these alterations provides a powerful tool for assessing tendon health and disease progression at a molecular level. Notably, the characteristic crosslinking profiles we identify could serve as functional biomarkers of tissue stiffness and viscoelasticity, factors central to tendon health and disease. The clear differences in crosslinking patterns between diseased and healthy tissues highlight the potential of these molecular signatures to differentiate between stages of tendon pathology and to inform treatment strategies aimed at modulating matrix composition and turnover.

The present study has several limitations which will need to be addressed in future research. One limitation of the human label-free proteomics analysis was the small number of patients and the use of gracilis tendon as healthy controls, rather than healthy rotator cuff tendons. A second limitation was the employment of a constitutive *Scx*^Cre^ driver instead of an inducible *Scx*^Cre^ERT2 system. Although an inducible system would have allowed more precise temporal control of *Vhl* deletion in adult tendons, the avascular nature of adult tendons compromised Cre recombination efficiency by limiting tamoxifen delivery. Additionally, genetically- and overload-induced murine models of tendinopathy are unlikely to fully recapitulate the human condition (^77^).

In conclusion, our data underscore the critical role of HIF1α as a driver of pathological adaptation to chronic overload. These findings pave the way for novel therapeutic approaches targeting HIF1α signaling in early-stage tendinopathy and downstream HIF1α targets in chronic tendinopathy, potentially offering new avenues for clinical management of tendon disorders.

## MATERIALS AND METHODS

### Study design

The objective of this study was to investigate the roles of mechanical overload and HIF1α activation in tendinopathy to address gaps in mechanistic understanding and develop more effective therapeutic strategies. Initial evaluation of diseased human biceps tendons showed hypoxic signaling and extensive ECM remodeling. Using genetically modified mouse models with targeted deletions of VHL, VEGFA, and/or HIF1α in tendon cells, we found that HIF1α stabilization and activation alone drives tendinopathy and disrupts the tendon’s load-bearing core, independent of blood vessels. Additionally, *in vitro* and *in vivo* models demonstrated that mechanical overload robustly activates HIF1α in tendon fibroblasts, and loss of HIF1α significantly improves overload-induced tendon maladaptations. For the *in vivo* overload model, mice were randomly assigned to one of the groups. The sample size for *in vivo* and *in vitro* experiments were estimated based on prior experience with the experimental models. Sample size and replicates are provided in figure legends.

## Study approval

Human study procedures and protocols were approved by the institutional review board of the Canton of Zurich (permissions Nr: 2013-0352, 2016-02665, 2020– 01119). Full informed consent was obtained from all patients. Animal studies were approved by the local committees on animal research (Kantonales Veterinäramt Zurich ZH083/2020 and ZH065/2023).

## Human samples

Long head biceps (LHB) tendons were harvested by subpectoral biceps tenodesis from symptomatic patients undergoing shoulder surgery (N=28; age: 48.36 ± 10 years). Rotator cuff tear tendon (subscapularis) biopsies were collected at the time of surgical debridement of the edges of the torn tendons (N=4, age: 60.5 ± 9 years). All patients were symptomatic and had full-thickness tendon tears on MRI. Healthy gracilis tendons were collected from patients undergoing surgical reconstruction of their anterior cruciate ligament or medial patellofemoral ligament (N= 17; age: 35.52 ± 12 years). Exclusion criteria for all patients in this study included other knee or shoulder pathology, rheumatoid arthritis, and systemic inflammatory disease.

## Mice

Mice were housed fed under specific pathogen-free conditions and received food and water ad libitum according to Swiss guidelines. Mouse strains were made in house or sourced from Jackson Laboratory or from other investigators (**Supplementary Table S2**). Data from mixed-sex groups were used, except for the B6/J experiments were only male were included to reduce variability. All the mouse strains were used at 7-8 weeks of age. Cre-negative mice from all the different mouse strains are named wild type (WT).

## Synergist Ablation

Adult mice received one injection of Temgesic (Buprenorphine; 0.1mg/kg s.c.) as preoperative analgesia 30 minutes before surgery. Bilateral Achilles tenectomy was performed under 2% isoflurane by transecting the Achilles tendon as previously described (^52^). After surgical intervention, the incision site was sutured with 8/O prolene suture (Ethicon, W8703) and mice received a second injection of Temgesic (Buprenorphine; 0.1mg/kg s.c.). Mice were euthanized with CO2 after 12 weeks post-surgery. Plantaris and Achilles tendon/ neotendon from both hindlegs were collected for histological analysis/immunofluorescence or mechanical test and crosslink analysis.

## In vitro stretching model

Tendon cells were isolated from gracilis human tendons and expanded in DMEM Low Glucose (Gibco, 31885023) with 10% heat inactivated fetal bovine serum (FBS, Gibco, 10500) and 1% P/S at 37°C and 5% CO2 until 80-90% confluency. Cells were seeded on collagen-coated silicon chambers (SYLGARD, Biesterfeld, 5498840000; curated at 40°C) at a density of 100.000 cells per chamber. After 3 days of culture with 10% FBS and 2 mM ascorbic acid (Wako, 1713265-25-8), the medium was replaced with 1% FBS without ascorbic acid. On the following day, the cells were cyclically stretched using a custom-made bioreactor at a frequency of 5%/s.

## Histology

For histology experiments, human and mouse tendon samples were harvested and immediately frozen in tissue freezing medium (Leica Biosystems). Sections (10-15μm thick) were cut using a cryostat (Leica) equipped with a low-profile blade (BioSys Laboratories, DB80 LX) and stored at -80°C until further use. Pathological phenotypes were identified with hematoxylin and eosin (H&E) and Picrosirius red (PSR, Polysciences, 24901) staining. Coded labelled images were evaluated and scored by the pathologist using the modified Bonar scoring system (^78,79^). H&E-stained sections were imaged using a Nikon-Eclipse-T2 objective with a 10x/0.45 objective. Picrosirius red-stained sections were imaged under polarized light using a Zeiss AxioImager.Z2 (ZEISS) equipped with a 20x/0.5 EC Epiplan-NeoFluar HD DIC objective. H&E and PSR image analysis was performed with ImageJ. Collagen fiber quantification was performed using the following threshold settings: Red (2-9/200-255), Orange (^10–39^), Yellow (^39–51^), Green (^52–128^).

## Immunofluorescence staining

Both histological slides as well as cells seeded on a silicone chamber were fixed with 4% paraformaldehyde (Carl Roth, 3105.2) for 10 min at room temperature (RT) and subsequently permeabilized and blocked with 3% BSA (Biowest, P6154) in PBST (PBS + 0.2% TritonX) for 1h at RT. Samples were then incubated overnight with primary antibody solution at 4°C and with secondary antibody for 1h at RT. Nuclei were counterstained with NucBlue (Thermo Fisher Scientific, R37606) for 20 minutes. After washing, cells were immersed in PBS while histological slides were mounted on a glass coverslip with mounting medium (Fisher Scientific, 9990412). Antibodies list can be found in **Supplementary Table S4**. Human tissue images were acquired with a ZEISS Axioscan 7 slide scanner (ZEISS) equipped with a 20x/0.8 air objective, except for the HIF1α nuclear staining, which was imaged using an Olympus FluoView FV3000 (Olympus Life Science) confocal microscope with 30x/1.05 silicon objective. Tenocyte images were acquired with a Leica TCS SP8 (Leica) confocal microscope using a 10x/0.3 air objective.

## Whole-mount staining

For 3D-whole mount imaging of *ScxGFP+* cells, Achilles tendon was fixed with 4% paraformaldehyde (Carl Roth, 3105.2) for 1.5 hours and placed between a glass slide and a cover slip with PBS. Images were acquired with Olympus FV3000 (Olympus Life Science) confocal microscope using a 60x/1.35 oil objective. 3D-whole mount tendon staining of blood vessels (CD31^+^) and myelinated nerve cells (NF-M+) was performed according to the SUMIC protocol (^80^). Cleared samples were imaged with a Leica TCS SP8 (Leica) confocal microscope and a 20x/0.75 immersion objective. 3D tissue reconstruction was performed with ImageJ.

## Generation of CRISPR/Cas9 -mediated VHL knockout cells

Four single guides RNAs (sgRNAs) against VHL gene were designed with CRISPOR online tool (http://crispor.tefor.net/) while a AAVS1 targeting sequence was used as control (**Supplementary Table S3**). Target sequence annealed oligos were cloned into the lentiCRISPRv2-puro transfer plasmid, a gift from Brett Stringer (Addgene plasmid #98290) (^81^) using a golden gate approach (VHL-targeting plasmids) or sequential digestion and ligation (AAVS1 control plasmid) using the Esp3I restriction enzyme (NEB, R0734S) and T4 DNA Ligase (NEB, M0202S). After lentiviral particle production, primary isolated human tendon cells were transduced 24 hours in the presence of 8 µg/ml polybrene and re-fed with fresh media the next day. Subsequently, human cells were selected with 3µg/ml puromycin (Gibco, A1113803) for one week. Knockout efficiency was verified by western blotting.

## Second Harmonic Generation (SHG)

Second harmonic generation of whole-mount Achilles tendons was performed as previously described (^51^). Image analysis was conducted with ImageJ by averaging the intensity signal over the planes and different locations for each sample. Collagen fibers dispersion was quantified using the ImageJ plugin ‘Directionality’.

## LOX-activatable fluorescent probe

The LOX-activatable fluorescent probe staining was performed as described previously (^82^). The fluorescent probe was imaged using an upright Leica TCS SP8 (Leica) two-photon microscope, equipped with a ×25/1 NA water-immersion objective at 760 nm excitation and 450-485 nm emission wavelengths. For each sample, *z* stacks were acquired with 15 planes at 0.5 um increment at two different locations. Image analysis was performed with ImageJ by averaging the intensity signal over the planes and different locations for each sample.

## Transmission electron microscopy

Tail tendon fascicles were freshly isolated from 7 weeks old mice in the morning (normal light/dark cycle), fixed and processed as described elsewhere (^51^). Segmentation of the cross-sectional area of collagen fibrils was performed with the Trainable Weka Segmentation ImageJ plugin (^83^).

## Calcium signaling

Calcium signaling response was investigated according to the protocol described in *Passini et al.* (^51^). Shortly, tenocytes seeded at a density of 100.000 cells/chamber and cultured for 3 days were stained with a calcium staining mix (5μM fluo-4 AM ;Thermo Fisher Scientific, F14217) and NucBlue (Thermo Fisher Scientific, R37605) in a modified Krebs–Henseleit (KHS) solution for 1.5 hours at 37°C and 5% O2. On the other end, single tail tendon fascicles were stained for 2 hours at 29°C and 3% O2. Cells were stretched at 0.5% strain per second until 50% strain. Whereas single tendon fascicles were preloaded to 0.02 N (0% strain) and stretched at 0.25% strain per second until failure. Images were acquired using an iMic widefield microscope (Thermo Fisher Scientific) and cellular Ca^2+^ responses were analyzed using a custom script. Cells that showed an increase in Ca^2+^ Δ*F*/*F* > 1.5 within 30 s from the start of the stretching were categorized as active.

## Mechanical testing

The biomechanical properties of tail tendon fascicles (TTFs) and plantaris tendons were tested using a uniaxial test device (Zwick Z010 TN, 20 N load-cell). During testing, tendons were kept in a custom chamber filled with PBS. The diameter of each TTFs was measured under a microscope (20x objective, Motic AE2000) while the diameter of each plantaris was captured with a 0.28X telecentric lens (Edmund Optics) attached to a camera (Canon EOS M50) and measured with ImageJ. For the ramp-to-failure test, TTFs were preloaded to 0.1 N (L0 corresponding to 0% strain) and preconditioned 5 times to 1% strain (preload reapplied after every cycle).

Subsequently, samples were ramped to failure at a constant strain-rate of 1% strain/s. Plantaris tendons were preloaded to 0.2 N, preconditioned 5 times to 1% strain and ramped to failure at a rate of 1% strain/s. In the stress-relaxation test, TTFs were preconditioned 10 times to 1% strain and incrementally strained at 0.5%/s to 2%, 4%, 6%, 8% and 10% strain, holding at each increment for 120s. Results were analyzed using custom-written Matlab scripts (Matlab®, R2023a, MathWorks, Inc.).

### Flow cytometry and *ScxGFP*+ cell sorting

For FACS analysis, single cell suspensions obtained from WT and *Scx*^Cre^;*Vhl*^f/f^ TTFs were incubated with CD31-PE and CD45-APC/fire 750 antibodies (**Supplementary Table S4**) for 45 min at 4°C with shaking while doublets and dead cells were excluded using SITOX Blue staining. For cell sorting, GFP+ CD31/D45 – cells from *ScxGFP+* - WT and -*Scx*^Cre^;*Vhl*^f/f^ mice were sorted using a 100 µm sorting chip into cell culture medium and plated into a 24 well plate. Sorting efficacy was confirmed by visualizing live GFP+ fluorescent cells after plating. All samples were analyzed and sorted using a Sony SH800S Cell Sorter (Sony Biotechnology Inc.). Data analysis was performed using FlowJo Version 10.6.1.

## Spheroid capillary sprouting assay

HUVECS spheroids were prepared as described elsewhere (^84^). Conditioned media from control and VHL-knockout tendon cells were added on top of the gel to induce sprouting. After 24 hours, spheroids were fixed with 4% PFA at RT and imaged with a Leica DM IL LED (Leica) microscope. Data analysis was performed using ImageJ.

## Angiogenic cytokines quantification

Conditioned medium was prepared from 80 to 90% confluent *ScxGFP+* cells sorted from WT and *Scx*^Cre^;*Vhl*^f/f^ TTFs and human primary WT and *VHL-KO* tendon cells. Angiogenic mouse cytokines were assessed with a Proteome Profiler Angiogenesis Array Kit (R&D Systems, ARY015) following the manufacturer’s guidelines. Membranes were revealed using a Clarity Max Western ECL Substrate (Bio-Rad Laboratories, 1705062) on ChemiDoc Imaging system (Bio-Rad Laboratories) and quantified using ImageJ. VEGFA measurements from mouse and human conditioned media were performed with U-PLEX Mouse VEGFA Assay (K152UVK-1) and U-PLEX Human VEGFA Assay (K151UVK-1), according to the manufacturer’s instructions (Meso Scale Discovery). Plates were read on a QuickPlex SQ120 Imager (Meso Scale Discovery) and electroluminescent data were analyzed with MSD Discovery Workbench software.

## TransAM HIF1α ELISA

Cell nuclear and cytoplasmic fractions were extracted using a nuclear extraction kit (Active Motif, 40010) as described in the manufacturer’s instructions. HIF1α activity was measured in 3µg of nuclear extract with the TransAM® HIF-1 kit (Active Motif, 47096) following the manufacturer’s instructions.

## Glycolysis

The glycolytic flux was investigated according to the protocol described in *Zhang et al.* (^85^).Briefly, cells were incubated for 2 hours in growth medium containing 0.4 mCi/ml [5-3H]-D-glucose (PerkinElmer). Thereafter, supernatant was transferred into glass vials sealed with rubber stoppers. 3H2O was captured in hanging wells containing a Whatman paper soaked with H2O over a period of 48 hours at 37°C to reach saturation. Radioactivity was determined in the paper by liquid scintillation counting.

## RNA isolation from tissues and cells and quantitative real-time PCR

RNA extraction from mouse Achilles tendon and TTFs was performed as previously described (^86^) while for cell lysates the PureLink RNA Mini kit (Invitrogen, 12183018A) was used following the manufacturer’s instructions. Isolated RNA was stored at -80°C or processed immediately for cDNA synthesis using a High-Capacity cDNA Reverse Transcription Kit (Applied Biosystems, 4374966) as described in the manufacturer’s instructions. Gene expression analysis was performed by quantitative real-time PCR with cDNA corresponding to 15 ng of starting RNA using Platinum™ II Hot-Start Green PCR Master Mix (Thermo Fisher Scientific, 14001013) for TaqMan probes or PowerUp SYBR Green Master Mix (Thermo Fisher Scientific, A25742) for custom-designed primers (**Supplementary Table S5**). Samples were amplified using a CFX Opus 384 Real-Time PCR System (Bio-Rad Laboratories). All experiments were run with technical duplicates. Relative gene expression levels were quantified using the 2–ddCT method with β2m and Anxa5 as reference genes.

## Next-generation RNA sequencing (RNA-Seq) and bioinformatics analysis

Library preparation and sequencing was performed by GENEWIZ® (Leipzig, Germany). In short, RNA integrity and yield were assessed with Agilent Fragment Analyzer (Agilent Technologies, USA). Total RNA was used to prepare the RNA-seq libraries with ribosomal depletion enrichment. Libraries were then sequenced on a NovaSeq 6000 sequencer (Illumina Inc, USA) and sequenced data were processed using Kallisto to generate a count file matrix for each individual sample. Samples were pooled together on a single dataset for downstream analysis and genes with one count or less across all samples were filtered out. Following DESeq2 analysis workflow (^87^), differential expression analysis was performed on the raw counts after estimation of size factors (controlling for differences in the sequencing depth of the samples), the estimation of dispersion values for each gene and fitting a generalized linear model. Pathways analysis of DEGS was performed using EnrichR (^88^) while gene set enrichment analysis (GSEA) was performed with the fgsea R package (^89^).

## Western Blotting

Human tenocytes were lysed directly with 80ul of 1X Laemmli buffer (Thermo Fisher Scientific, 15493939), while mouse tail tendon fascicles were minced in 100ul RIPA buffer (Sigma-Aldrich, R0278) supplemented with protease inhibitors (Abcam, ab270055). Total protein was quantified using precision red advance protein assay (Cytoskeleton, ADV02). Samples were then boiled for 5 min at 95°C for protein denaturation. Equal amounts of protein were loaded onto a 4-15% mini-PROTEAN TGX strain-free protein gel (Bio-Rad, 4568086 and 4568084). Gels were transferred onto polyvinylidene difluoride (PVDF) membranes using the Trans-Blot-Turbo Transfer System (Bio-Rad). Membranes were blocked with 5% non-fat dry milk/TBS- T for 1 h at room temperature and analyzed with immunoblot. Images were taken using UltraScence Pico Ultra Western Substrate (GeneDireX, CCH345-B) and the ChemiDoc MP imaging system (Bio-Rad). Antibodies used for immunoblots are listed in **Supplementary Table S4**.

## Mass- spectroscopy

### Collagen and Crosslinks analysis

Collagen peptide analysis of snap frozen human long head biceps and gracilis tendons was performed as previously described (^90^). Human tendon samples, together with mouse TTFs and plantaris tendons were processed for LC-MS/MS crosslink analysis as previously reported (^47^).

### Label- free proteomics

#### Human Tissue peptide extraction

Proteins were extracted by pressure cycling technology (60 cycles of 450000psi for 20 s, atmospheric pressure for 10 s, 33°C) from ca. 4mg tissue in a lysis buffer containing 4M Guanidine hydrochloride (GuHCl; Sigma-Aldrich, 50940) and 250mM HEPES, pH 7.8 (Sigma-Aldrich, 54457) using a Barocycler 2320ETX (Pressure Biosciences Inc.) followed by sonication at 4°C (10cycles: 30sec on/30sec off). Proteins were then reduced with 200mM tris(2-carboxyethyl) phosphine hydrochloride (TCEP; Thermo Fisher Scientific, PG82080) during 1h of shaking at 37°C and alkylated with 400mM chloroacetamide (CAA, Sigma-Aldrich, C0267) during 30min of shaking at room temperature. After diluting the GuHCl concentration to 1.2M with HEPES, 10µg of proteins were pre-digested in LysC (Promega, VA1170; 1:50 enzyme-to-protein ratio) for 4 hours at 37°C. Trypsin (Promega, V5111) digestion (1:20 enzyme-to-protein ratio) was performed overnight at 37°C and in 0.8M GuHCl.

#### Mouse tendon fascicles (TFFs) peptide extraction

Proteins were extracted by pressure cycling technology (60 cycles of 450000psi for 20 s, atmospheric pressure for 10 s, 33°C) from ca. 4mg tissue in a lysis buffer containing 4M Guanidine hydrochloride (GuHCl; Sigma-Aldrich, 50940) and 250mM HEPES, pH 7.8 (Sigma-Aldrich, 54457) using a Barocycler 2320ETX (Pressure Biosciences Inc.). DNA was digested was performed by adding 1.25U of Benzonase to each sample followed by 15 minutes incubation at RT. Proteins were then reduced with TCEP (10 mM final concentration) and alkylated with Iodoacetamide (IAA; Sigma-Aldrich, I1149; 40 mM final concentration) during 30 min of shaking at room temperature. After diluting the GuHCl concentration to 1.2M with HEPES, LysC (Promega, VA1170) was then added (1:100 enzyme-to-protein ratio), followed by a second barocycler step (45 cycles of 50s 20000psi and 10s atmospheric pressure at 33°C). Trypsin (Promega, V5111) digestion (1:50 enzyme-to-protein ratio) was performed with another barocycler step (90 cycles of 20000psi for 50s, atmospheric pressure for 10s at 33°C) and carried out overnight at 37°C.

#### Peptides cleanup, reverse phase liquid chromatography and acquisition of mass spectra

Peptides were acidified to pH 2 using 10% trifluoroacetic acid to stop the digestion and desalted on C18 cartridges (SepPak 50mg, Waters). Cartridges were preconditioned with methanol, activated with buffer B (80% acetonitrile, 0.1% formic acid, 19.9% water), and equilibrated twice with buffer A* (3% acetonitrile, 1% trifluoroacetic acid, 96% water) before loading the peptide samples. Peptides were washed four times with buffer A (0.1% acetonitrile, 99.9% water), eluted in buffer B, dried in a SpeedVac at 60°C (Eppendorf) and resuspended in MS buffer (2% acetonitrile, 1%trifluoroacetic acid, 97% water) containing iRTs (Biognosys). 500ng of human and mouse peptide mixtures were loaded onto 50cm x 75µm, PepMap™RSLC analytical column packed with 2µm C18 beads (Thermo Fisher Scientific, ES803A) and separated on a 2cm x 75µm, Acclaim®PepMap100 trap column, packed with 3µm, C18 beads (Thermo Fisher Scientific, 164946) using an EASY-nLC 1200 liquid chromatography system (Thermo Scientific Scientific) coupled in line with a QExactive mass spectrometer (Thermo Fisher Scientific). The peptides were eluting from the column at a mixture of solvent A (0.1% formic acid/99.9% water (Fisher Scientific, LS118-212) and solvent B (80% Acetonitrile, 0.1% Formic Acid, 19.9% water (Fisher Scientific, 15431423)) at a constant flow rate of 250nl/min. The chromatographic gradient ran from 10% to 95% solvent B in 14min (10% to 23% in 85 min, 23% to 38% in 30 min, 38% to 60% in 10 min followed by wash steps from 60% to 95% in 5min and 95% Solvent B for 10min). For mouse peptides an Orbitrap Fusion mass spectrometer (Thermo Fisher Scientific) was used for analysing. Peptide precursor m/z measurements were carried out at resolution of 70000 after accumulation to an automatic gain control (AGC) target of 3e6. Maximum injection time was set to 20ms, the scan range was between 300 and 1750m/z, selecting the top 10 MS1 ions for MS2 analysis. Higher-energy collisional dissociation MS2 scans were recorded using 17500 resolution for human and 35000 for mouse, applying an AGC target between 2e4 and 1e6 and a maximum injection time of 60ms (human) or 120ms (mouse) . Precursors were isolated using a window of 1.6 m/z and fragmented with a normalized collision energy (NCE) of 28% (human) or 35% (mouse).

#### Data analysis

Human and mouse raw data were analyzed using MaxQuant v1.6.17 software using the integrated Andromeda search engine. The database was concatenated with one composed of all protein sequences in reversed order. Methionine and proline oxidation, as well as protein N-terminal acetylation ere set as variable modifications. Peptide-spectrum matches (PSM’s) did not exceed 1% false discovery rate (FDR).

Filtered PSM’s were further filtered for peptide and protein-level FDR of 1%. To determine differentially abundant proteins between diseased and control human tendons, global bioinformatics analysis was conducted on the normalized label-free intensity values. Normalized counts were imported into Omics Playground Suite (BigOmics Analytics, Switzerland) implemented within RStudio (^91^). trend.limma algorithm was used to estimate the log2(FC) and FDR (meta.q) of differentially abundant proteins between the two groups (^92–94^). Significance was assumed at an FDR of 0.05 -log10(meta.q) >1.3. Heatmap hierarchical clustering was performed at the feature level (protein). Functional annotation of clusters (M1 and M2) was quired against reference database Molecular Signatures Database (MSigDB). Annotations were ranked based on the highest correlation (R) with the reference set. Top differentially enriched sets were determined with Gene Set Enrichment Analysis (GSEA) package (^95^). Volcano plots were visualized using VolcaNoseR package (^96^). Label-free quantification and differential expression analysis of mouse proteins were performed using the DEP analysis workflow (Bioconductor, 10.18129/B9.bioc.DEP) and the limma R package (^97^).

## Statistical analyses

Data are presented as mean ± SE, unless indicated otherwise. Group sizes were determined based on the results of preliminary experiments. Mice were assigned at random to groups. Experiments were not performed in a blinded fashion. Statistical significance for each experiment was determined as shown in the figure legends, where n= the number of independent biological replicates (humans and animals, unless noted as cells) per group. The age and sex of the human samples were tested as covariates. Statistical analyses were performed in Prism (GraphPad) or R studio. No data points were excluded.

## List of Supplementary Materials

Supplementary Materials and Methods Figs. S1 to S12

Tables S1 to S5

Data files S1 to S2

## Acknowledgments

The authors thank the donors who generously consented to donate their tendon tissue for research; clinical teams and research nurses, the administrative staff at Balgrist University Hospital, the Swiss Center for Musculoskeletal Biobanking and Schwerzenbach; C. Leuzinger for her dedicated care of the mice used in this study; P. Gilardoni and the Hiwis team for genotyping; members of the Snedeker group, De Bock group and Surdez group for constructive discussions; R. Schweitzer for providing *ScxCre* and *ScxGFP* mice; C. Stockmann for providing *Vhl^f/f^* and *Vegfa^f/f^* mice. The authors gratefully acknowledge the Functional Genomics Center Zurich (FGCZ) of University of Zurich and ETH Zurich, and in particular L. Opitz and S. Pfammatter, for the support on Genomics and Proteomics analyses. Imaging was performed with support of the Center for Microscopy and Image Analysis, University of Zurich. Schematics were made using Biorender.

## Author Contributions

Conceptualization: G.M., U.B., K.D.B. and J.G.S. Methodology: G.M., A.G.M., I.S.N., D.E.T., J.C., P.K.J., S.L.W., O.B., S.M., L.M., D.S., M.R.A., H.W. Investigation: G.M., A.G.M, I.S.N, D.E.T, J.C., P.K.J., S.L.W., O.B., B.N., M.B., M.W.,R.A., E.M. Formal analysis: P.K.J.,A.A.H., S.A.,G.T., M.H. Resources: F.S., K.W., S.F.F., M.R.A., H.W. Software: P.K.J., S.A., G.T. Visualization: G.M., A.A.H. Funding acquisition: U.B., D.E., K.D.B and J.G.S. Project administration: K.D.B and J.G.S. Supervision: A.A.H., F.S.P., K.D.B. and J.G.S. Writing – Original Draft Preparation: G.M., F.S.P., K.D.B. and J.G.S.

## Data and materials availability

All data associated with this study are in the paper and its Supplementary Materials. Mouse raw and processed sequencing data are available at GSE276425. Mouse proteomics and human transcriptomics /proteomics raw data are available upon request.

