## Supplementary Materials and Methods Figs. S1 to S12 Tables S1 to S5 Data files S1 to S2 for "HIF1α gates tendon response to overload and drives tendinopathy independently of vascular recruitment"

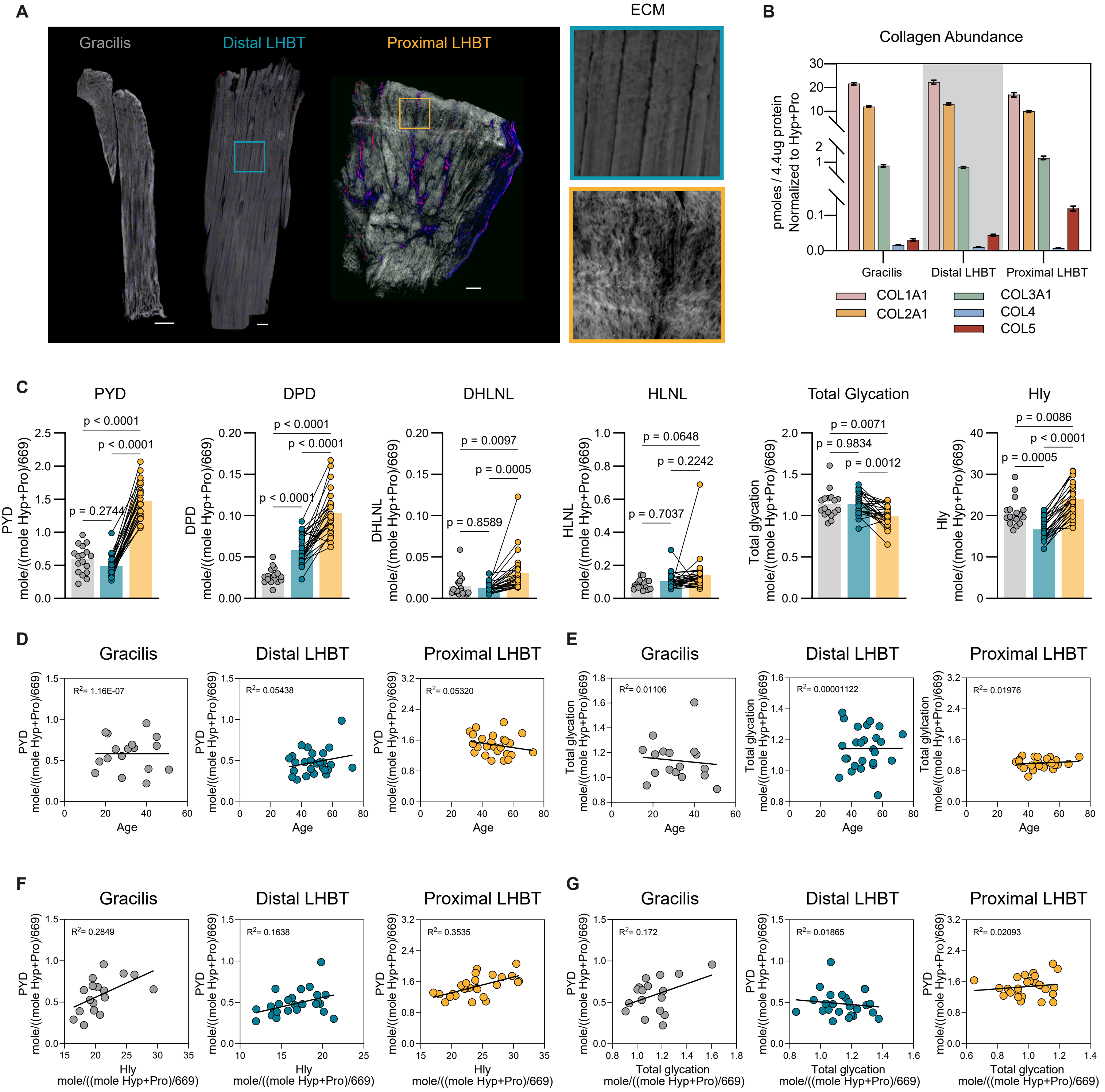

A
Gracilis
Distal LHBT
Proximal LHBT
ECM
B
Collagen Abundance
pmoles / 4.4ug protein Normalized to Hyp+Pro
Gracilis
Distal LHBT
Proximal LHBT
COL1A1
COL2A1
COL3A1
COL4
COL5
C
PYD
DPD
DHLNL
HLNL
Total Glycation
Hly
mole/((mole Hyp+Pro)/669)
p < 0.0001
p = 0.2744
p = 0.0097
p = 0.0005
p = 0.8589
p = 0.0648
p = 0.2242
p = 0.7037
p = 0.0071
p = 0.9834
p = 0.0012
p = 0.0086
p = 0.0005
D
E
Age
Total glycation
$R^2$= 1.16E-07
$R^2$= 0.05438
$R^2$= 0.05320
$R^2$= 0.01106
$R^2$= 0.00001122
$R^2$= 0.01976
F
G
$R^2$= 0.2849
$R^2$= 0.1638
$R^2$= 0.3535
$R^2$= 0.172
$R^2$= 0.01865
$R^2$= 0.02093

**Fig. S1. Human diseased tendons display matrix dysregulation and aberrant collagen crosslinks.** (**A**) Representative images of human healthy gracilis (n=17) and diseased long head biceps tendons (n=28) stained for the endothelial cell marker (CD31, red) and cell nuclei (NucBlue, cyan). Autofluorescence signal detected in the 488 channel was used to visualize ECM (grey). Scale bar: 100 µm. (**B**) Mass-spectrometry based peptide quantification of COL1A1, COL2A1, COL3A1, COL4 and COL5 in gracilis (n=17) and long head biceps tendons (n=26). Collagen peptides are expressed in picomoles per 4.4 microgram total protein normalized to the sum of hydroxyproline and proline (Hyp+Pro) content measured by amino acid analysis. Bar graphs represent mean ± SEM. (**C**) Enzymatic crosslinks, total glycation and hydroxylysine (Hly) levels normalized by the sample Hyp+Pro content and divided by total number of Hyp+Pro residues in collagen I (699) in human gracilis (n=17) and long head biceps tendons (n=26). Data are shown as mean with individual points representing biologically independent samples and lines connecting paired samples. One-way ANOVA with Tukey's multiple comparisons test was used for statistical analysis (**D**) Linear regression analysis shows no correlation between pyridinoline (PYD) levels and patient age and (**E**) total glycation levels and patient age in both human gracilis (n=17) and long head biceps tendons (n=26). (**F**) Linear regression analysis shows a positive correlation between pyridinoline (PYD) levels and hydroxylysine (Hly) content in both human gracilis (n=17) and long head biceps tendons (n=26). (**G**) Linear regression analysis shows a positive correlation between pyridinoline (PYD) levels and total glycation in human gracilis (n=17) while no strong correlation is present in the long head biceps tendons (n=26). Each point represents a biologically independent samples in D,E,F,G.

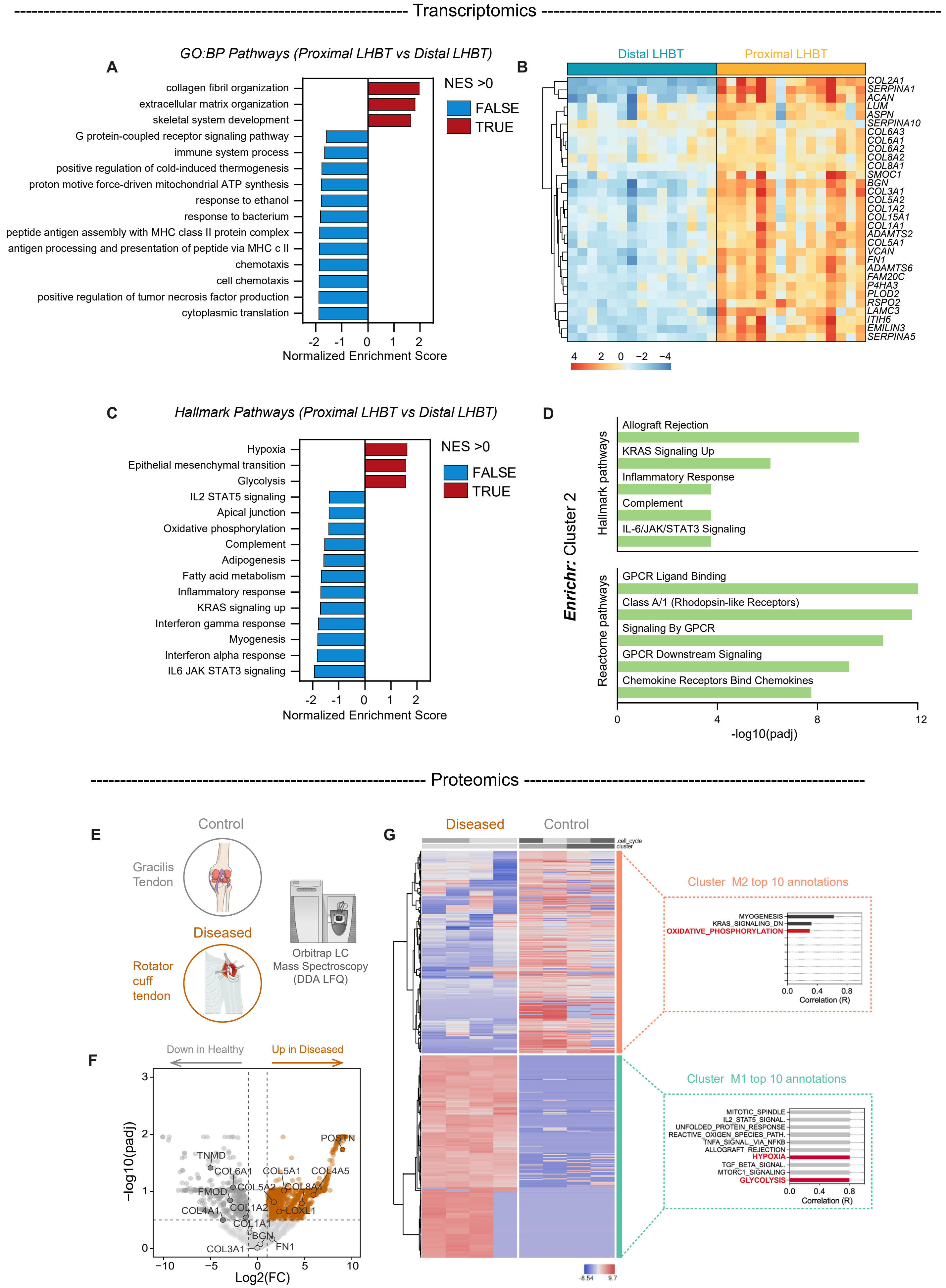
Transcriptomics
A
GO:BP Pathways (Proximal LHBT vs Distal LHBT)
collagen fibril organization
extracellular matrix organization
skeletal system development
G protein-coupled receptor signaling pathway
immune system process
positive regulation of cold-induced thermogenesis
proton motive force-driven mitochondrial ATP synthesis
response to ethanol
response to bacterium
peptide antigen assembly with MHC class II protein complex
antigen processing and presentation of peptide via MHC c II
chemotaxis
cell chemotaxis
positive regulation of tumor necrosis factor production
cytoplasmic translation
Normalized Enrichment Score
NES >0
FALSE
TRUE
B
Distal LHBT
Proximal LHBT
COL2A1
SERPINA1
ACAN
LUM
ASPN
SERPINA10
COL6A3
COL6A1
COL6A2
COL8A2
COL8A1
SMOC1
BGN
COL3A1
COL5A2
COL1A2
COL15A1
COL1A1
ADAMTS2
COL5A1
VCAN
FN1
ADAMTS6
FAM20C
P4HA3
PLOD2
RSPO2
LAMC3
ITIH6
EMILIN3
SERPINA5
C
Hallmark Pathways (Proximal LHBT vs Distal LHBT)
Hypoxia
Epithelial mesenchymal transition
Glycolysis
IL2 STAT5 signaling
Apical junction
Oxidative phosphorylation
Complement
Adipogenesis
Fatty acid metabolism
Inflammatory response
KRAS signaling up
Interferon gamma response
Myogenesis
Interferon alpha response
IL6 JAK STAT3 signaling
Normalized Enrichment Score
NES >0
FALSE
TRUE
D
Enrichr: Cluster 2
Hallmark pathways
Allograft Rejection
KRAS Signaling Up
Inflammatory Response
Complement
IL-6/JAK/STAT3 Signaling
Reactome pathways
GPCR Ligand Binding
Class A/1 (Rhodopsin-like Receptors)
Signaling By GPCR
GPCR Downstream Signaling
Chemokine Receptors Bind Chemokines
-log10(padj)
Proteomics
E
Control
Gracilis Tendon
Diseased
Rotator cuff tendon
Orbitrap LC Mass Spectroscopy (DDA LFQ)
F
Down in Healthy
Up in Diseased
-log10(padj)
Log2(FC)
TNMD
COL6A1
COL5A1
COL4A5
POSTN
FMOD
COL5A2
COL8A1
COL4A1
COL1A2
LOXL1
COL1A1
BGN
FN1
COL3A1
G
Diseased
Control
Cluster M2 top 10 annotations
MYOGENESIS
KRAS_SIGNALING_DN
OXIDATIVE_PHOSPHORYLATION
Correlation (R)
Cluster M1 top 10 annotations
MITOTIC_SPINDLE
IL2_STAT5_SIGNAL.
UNFOLDED_PROTEIN_RESPONSE
REACTIVE_OXIGEN_SPECIES_PATH.
TNFA_SIGNAL._VIA_NFKB
ALLOGRAFT_REJECTION
HYPOXIA
TGF_BETA_SIGNAL.
MTORC1_SIGNALING
GLYCOLYSIS
Correlation (R)
-8.54
9.7

**Fig. S2. Human pathological tendons exhibit enhanced matrix turnover and a metabolic switch at the transcriptomic level.** (**A**) Bar plots showing Normalized Enrichment Score of significant (adjusted p-value $< 0.05$) biological processes pathways (Gene Ontology) from GSEA analysis conducted on proximal LHBT samples versus distal LHBT control (n=15). (**B**) Heatmap of centered normalized counts showing ECM-related genes selected from Matrisome DB2.0 database. Columns represent individual samples (n=15). (**C**) Bar plots showing Normalized Enrichment Score of significant (padj $< 0.05$) Hallmark pathways (MsigDB) from GSEA analysis conducted on proximal LHBT samples versus distal LHBT control (n=15). (**D**) Enriched pathway analysis of Cluster 2 using *Enrichr* queried against MSigDB Hallmark and Reactome databases. (**E**) Schematic representation of anatomical sourcing from two human cohorts used for label-free proteomics analysis. Healthy gracilis tendons (n=4) were harvested as graft material from patients undergoing anterior cruciate ligament (ACL) or medial patellofemoral ligament (MPFL) reconstruction while diseased rotator cuff (RCT)tendons (supraspinatus, n=4) biopsies were harvested from patients undergoing repair shoulder arthroscopy. (**F**) Volcano plot of differentially expressed genes (DEGs) in diseased RCT group relative to the gracilis control group. Colored dots show the 1624 proteins corresponding to a cutoff of -log10(padj>0.5) and log2FC±1. (**G**) Heatmap of protein-level hierarchical clustering of the differentially abundant proteins in healthy vs. diseased tendons. Columns represent individual samples (n=4). Blue denotes less-abundant proteins; red denotes more-abundant proteins. Bar plots indicate the fisher-weighted, average correlations of each cluster with the MSigDB Hallmark database annotation terms. Differentially abundant protein shows positive correlation with hypoxia and glycolysis signatures in the diseased group while negatively-correlates with oxidative phosphorylation signature.

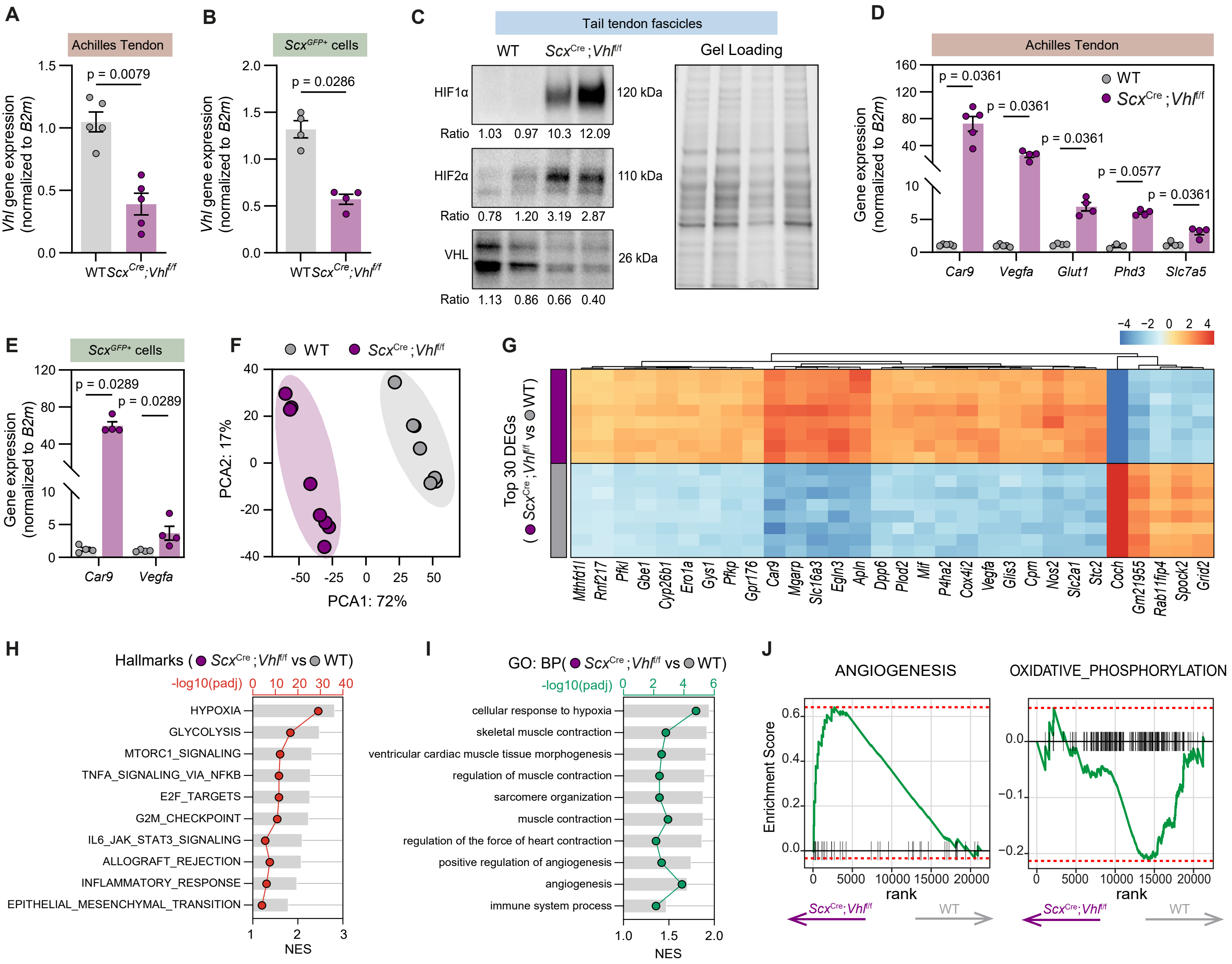
A
Achilles Tendon
p = 0.0079
Vhl gene expression (normalized to B2m)
WT
ScxCre;Vhlf/f
B
ScxGFP+ cells
p = 0.0286
Vhl gene expression (normalized to B2m)
WT
ScxCre;Vhlf/f
C
Tail tendon fascicles
WT
ScxCre;Vhlf/f
Gel Loading
HIF1α
120 kDa
Ratio 1.03 0.97 10.3 12.09
HIF2α
110 kDa
Ratio 0.78 1.20 3.19 2.87
VHL
26 kDa
Ratio 1.13 0.86 0.66 0.40
D
Achilles Tendon
WT
ScxCre;Vhlf/f
p = 0.0361
p = 0.0361
p = 0.0361
p = 0.0577
p = 0.0361
Gene expression (normalized to B2m)
Car9
Vegfa
Glut1
Phd3
Slc7a5
E
ScxGFP+ cells
p = 0.0289
p = 0.0289
Gene expression (normalized to B2m)
Car9
Vegfa
F
WT
ScxCre;Vhlf/f
PCA2: 17%
PCA1: 72%
G
Top 30 DEGs (ScxCre;Vhlf/f vs WT)
Mthfd1l
Rnf217
Pfkl
Gbe1
Cyp26b1
Ero1a
Gys1
Pfkp
Gpr176
Car9
Mgarp
Slc16a3
Egln3
Apln
Dpp6
Plod2
Mif
P4ha2
Cox4i2
Vegfa
Glis3
Cpm
Nos2
Slc2a1
Stc2
Coch
Gm21955
Rab11fip4
Spock2
Grid2
H
Hallmarks (ScxCre;Vhlf/f vs WT)
-log10(padj)
HYPOXIA
GLYCOLYSIS
MTORC1_SIGNALING
TNFA_SIGNALING_VIA_NFKB
E2F_TARGETS
G2M_CHECKPOINT
IL6_JAK_STAT3_SIGNALING
ALLOGRAFT_REJECTION
INFLAMMATORY_RESPONSE
EPITHELIAL_MESENCHYMAL_TRANSITION
NES
I
GO: BP(ScxCre;Vhlf/f vs WT)
-log10(padj)
cellular response to hypoxia
skeletal muscle contraction
ventricular cardiac muscle tissue morphogenesis
regulation of muscle contraction
sarcomere organization
muscle contraction
regulation of the force of heart contraction
positive regulation of angiogenesis
angiogenesis
immune system process
NES
J
ANGIOGENESIS
OXIDATIVE_PHOSPHORYLATION
Enrichment Score
rank
ScxCre;Vhlf/f
WT

**Fig. S3. *Vhl*-knockout leads to activation of HIF-dependent angiogenic and metabolic transcriptional program.** (**A**) *Vhl* mRNA levels in Achilles tendon (n=5) and (**B**) TTFs-isolated *Scx*$^{GFP+}$ tenocytes (n=4) from WT and *Scx*$^{Cre}$;*Vhl*$^{f/f}$ mice. (**C**) HIF1α, HIF2α and VHL protein levels normalized for total protein in WT and *Scx*$^{Cre}$;*Vhl*$^{f/f}$ TTFs (n=2). Densitometric quantification is shown below the lanes. (**D**) mRNA levels of canonical HIF1α target genes in Achilles tendon (n=5) and (**E**) TTFs (n=4) of WT and *Scx*$^{Cre}$;*Vhl*$^{f/f}$ mice. (**F**) Principal component analysis showing separation of WT and *Scx*$^{Cre}$;*Vhl*$^{f/f}$ TTFs at the transcriptome level (n=8). (**G**) Heatmap of centered normalized counts showing the top 30 DEGs in *Scx*$^{Cre}$;*Vhl*$^{f/f}$ compared to WT TTFs. Columns represent individual samples (n=8). (**H**) Bar plots showing Normalized Enrichment Score and dot plot showing the -log10(padj) values of significant (padj < 0.05) Hallmark pathways (MsigDB) and (**I**) Biological Processes pathways (Gene Ontology) from GSEA analysis. The analysis was conducted on *Scx*$^{Cre}$;*Vhl*$^{f/f}$ versus WT TTFs (n=8). (**J**) GSEA of angiogenesis and oxidative phosphorylation pathways conducted on transcriptomics data against Hallmark database (MsigDB) of *Scx*$^{Cre}$;*Vhl*$^{f/f}$ versus WT TTFs (n=8). Each point represents a single mouse in A,B,D,E,F. Bar graphs indicate mean ± SEM. Student's t test (unpaired, two tailed) was used in A,B. Multiple Mann-Whitney tests were used in D,E.

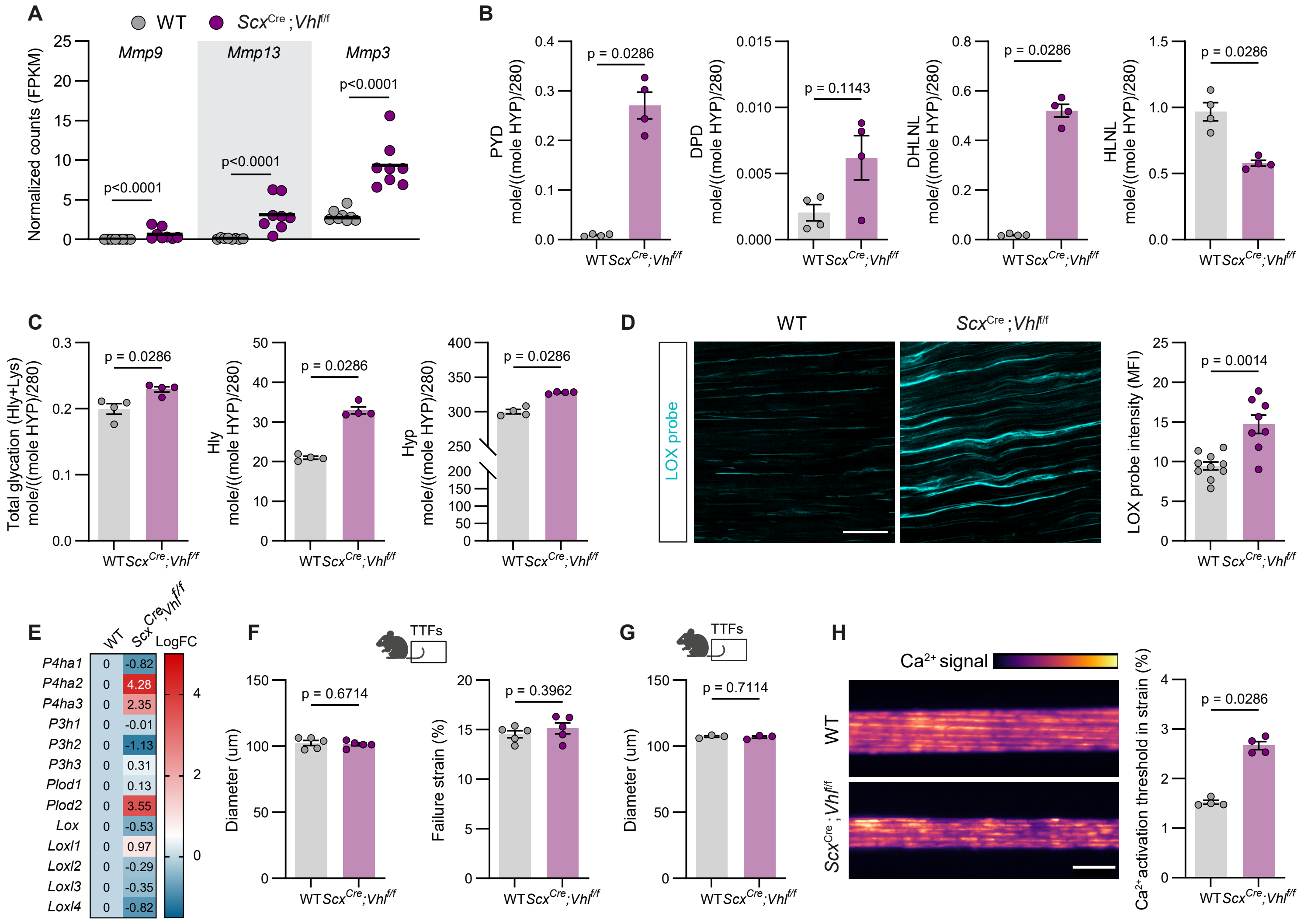
A
WT
ScxCre;Vhlf/f
Mmp9
Mmp13
Mmp3
Normalized counts (FPKM)
p<0.0001
p<0.0001
p<0.0001
B
PYD mole/((mole HYP)/280)
p = 0.0286
DPD mole/((mole HYP)/280)
p = 0.1143
DHLNL mole/((mole HYP)/280)
p = 0.0286
HLNL mole/((mole HYP)/280)
p = 0.0286
C
Total glycation (Hly+Lys) mole/((mole HYP)/280)
p = 0.0286
Hly mole/((mole HYP)/280)
p = 0.0286
Hyp mole/((mole HYP)/280)
p = 0.0286
D
WT
ScxCre;Vhlf/f
LOX probe
LOX probe intensity (MFI)
p = 0.0014
E
WT
ScxCre;Vhlf/f
LogFC
P4ha1 0 -0.82
P4ha2 0 4.28
P4ha3 0 2.35
P3h1 0 -0.01
P3h2 0 -1.13
P3h3 0 0.31
Plod1 0 0.13
Plod2 0 3.55
Lox 0 -0.53
Loxl1 0 0.97
Loxl2 0 -0.29
Loxl3 0 -0.35
Loxl4 0 -0.82
F
TTFs
Diameter (um)
p = 0.6714
Failure strain (%)
p = 0.3962
G
TTFs
Diameter (um)
p = 0.7114
H
Ca2+ signal
WT
ScxCre;Vhlf/f
Ca2+ activation threshold in strain (%)
p = 0.0286

**Fig. S4. *Scx*$^{Cre}$;*Vhl*$^{f/f}$ tendons show higher matrix turnover and collagen crosslinking which causes impaired mechanics and mechanotransduction**. (**A**) *Mmp9*, *Mmp13* and *Mmp3* mRNA levels expressed as normalized counts from bulk RNA-seq dataset of WT and *Scx*$^{Cre}$;*Vhl*$^{f/f}$ TTFs (n=8). (**B**) Mature, immature crosslinks levels and (**C**) total glycation level, hydroxylysine and hydroxyproline content normalized by Hyp content divided by total number of Hyp residues in collagen I (280) in WT and *Scx*$^{Cre}$;*Vhl*$^{f/f}$ TTFs (n=4). (**D**) Representative images (left panel) and quantification (right panel) of LOX activity by utilizing a fluorescent probe in WT (n=10) and *Scx*$^{Cre}$;*Vhl*$^{f/f}$ (n=8) Achilles cryosections. Scale bar: 50μm. (**E**) Collagen crosslinking enzyme mRNA levels in WT and *Scx*$^{Cre}$;*Vhl*$^{f/f}$ TTFs (n=8) expressed as Log2FC. (**F**) Diameter and failure strain values from WT and *Scx*$^{Cre}$;*Vhl*$^{f/f}$ TTFs (n=5) used in the ramp to failure mechanical test and (**G**) stress relaxation test. (**H**) Representative images (left panel) and quantification (right panel) of calcium signaling activation threshold from WT and *Scx*$^{Cre}$;*Vhl*$^{f/f}$ TTFs (n=4). Threshold is expressed in strain %. Scale bar: 50μm. Each point represents a single mouse. Bar graphs indicate mean ± SEM. Student's t test (unpaired, two tailed) was used in A,B,C,D,F,G,H.

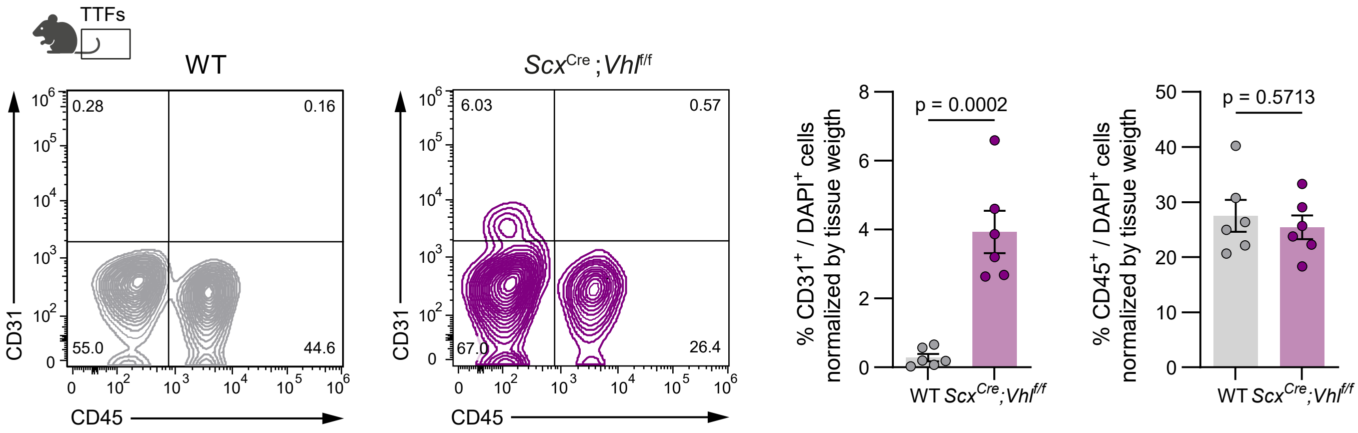

TTFs
WT
ScxCre;Vhlf/f
0.28
0.16
55.0
44.6
6.03
0.57
67.0
26.4
CD31
CD45
p = 0.0002
% CD31+ / DAPI+ cells normalized by tissue weigth
p = 0.5713
% CD45+ / DAPI+ cells normalized by tissue weigth
WT
ScxCre;Vhlf/f

**Fig. S5. Flow cytometry gating strategy for endothelial cells and immune cells in tail tendon fascicles cell suspension.** Flow cytometry gating strategy (left panels) and quantification (right panels) of $CD31^{+}$ ECs and $CD45^{+}CD31^{-}$ immune cells in the tail tendon fascicles of $\textit{Scx}^{Cre}$;$\textit{Vhl}^{f/f}$ TTFs and their wild type littermates (n=6). Each point represents a single mouse. Bar graphs indicate mean ± SEM. Student's t test (unpaired, two tailed) was used for statistical analysis.

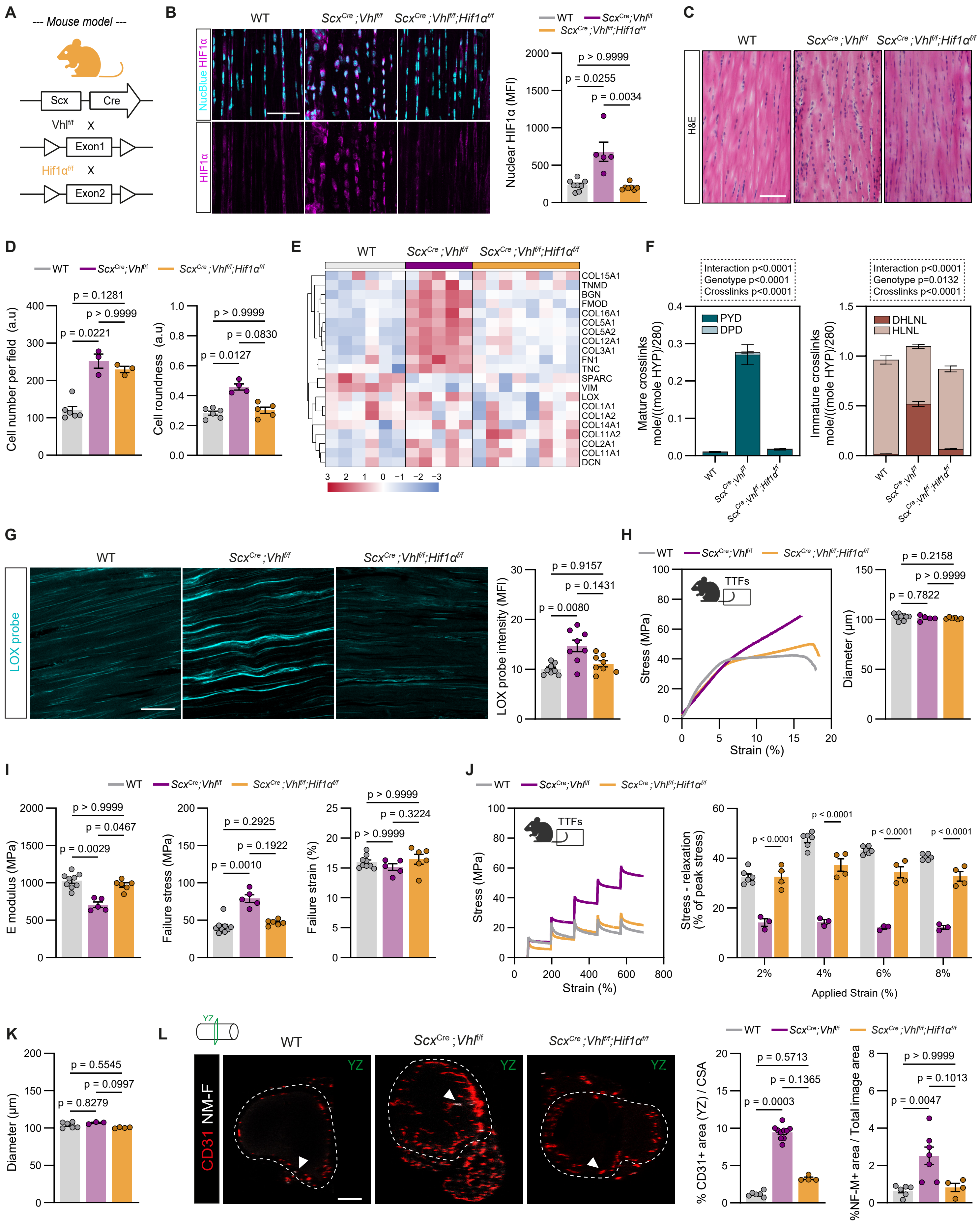

A
--- Mouse model ---
Scx
Cre
Vhl f/f X
Exon1
Hif1α f/f X
Exon2
B
WT
ScxCre;Vhlf/f
ScxCre;Vhlf/f;Hif1αf/f
NucBlue HIF1α
HIF1α
Nuclear HIF1α (MFI)
p > 0.9999
p = 0.0255
p = 0.0034
C
H&E
D
Cell number per field (a.u)
p = 0.1281
p > 0.9999
p = 0.0221
Cell roundness (a.u)
p > 0.9999
p = 0.0830
p = 0.0127
E
COL15A1
TNMD
BGN
FMOD
COL16A1
COL5A1
COL5A2
COL12A1
COL3A1
FN1
TNC
SPARC
VIM
LOX
COL1A1
COL1A2
COL14A1
COL11A2
COL2A1
COL11A1
DCN
F
Interaction p<0.0001
Genotype p<0.0001
Crosslinks p<0.0001
Mature crosslinks mole/((mole HYP)/280)
PYD
DPD
Interaction p<0.0001
Genotype p=0.0132
Crosslinks p<0.0001
Immature crosslinks mole/((mole HYP)/280)
DHLNL
HLNL
G
LOX probe
LOX probe intensity (MFI)
p = 0.9157
p = 0.1431
p = 0.0080
H
TTFs
Stress (MPa)
Strain (%)
Diameter (µm)
p = 0.2158
p > 0.9999
p = 0.7822
I
E modulus (MPa)
p > 0.9999
p = 0.0467
p = 0.0029
Failure stress (MPa)
p = 0.2925
p = 0.1922
p = 0.0010
Failure strain (%)
p > 0.9999
p = 0.3224
p > 0.9999
J
TTFs
Stress (MPa)
Strain (%)
Stress - relaxation (% of peak stress)
p < 0.0001
Applied Strain (%)
2%
4%
6%
8%
K
Diameter (µm)
p = 0.5545
p = 0.0997
p = 0.8279
L
YZ
CD31 NM-F
% CD31+ area (YZ) / CSA
p = 0.5713
p = 0.1365
p = 0.0003
%NF-M+ area / Total image area
p > 0.9999
p = 0.1013
p = 0.0047

**Fig. S6. The pathological tendon phenotype observed in *Scx*$^{Cre}$;*Vhl*$^{f/f}$ mice is HIF1α-dependent.** (**A**) Generation of *Scx*$^{Cre}$;*Vhl*$^{f/f}$;*Hif1α*$^{f/f}$ mice expressing constitutive Cre under the *Scx* promoter. (**B**) Representative images (left panel) and quantification (right panel) of HIF1α nuclear translocation based on colocalization with NucBlue staining in WT (n=8), *Scx*$^{Cre}$;*Vhl*$^{f/f}$ (n=5) and *Scx*$^{Cre}$;*Vhl*$^{f/f}$;*Hif1α*$^{f/f}$ (n=7) Achilles tendon. Scale bar: 50μm. (**C**) Representative images of haematoxylin-eosin (H&E)-stained sections and (**D**) cell number and cell roundness quantification in WT (n=6), *Scx*$^{Cre}$;*Vhl*$^{f/f}$ (n=3) and *Scx*$^{Cre}$;*Vhl*$^{f/f}$;*Hif1α*$^{f/f}$ (n=3) Achilles tendon. Scale bar: 100μm. (**E**) Heatmap of ECM-related proteins expressed as centered normalized intensities in WT (n=6), *Scx*$^{Cre}$;*Vhl*$^{f/f}$ (n=5) and *Scx*$^{Cre}$;*Vhl*$^{f/f}$;*Hif1α*$^{f/f}$ (n=8) TTFs. (**F**) Mature and immature crosslink levels normalized by Hyp content divided by total number of Hyp residues in collagen I (280) in WT (n=11), *Scx*$^{Cre}$;*Vhl*$^{f/f}$ (n=4) and *Scx*$^{Cre}$;*Vhl*$^{f/f}$;*Hif1α*$^{f/f}$ (n=6) TTFs. (**G**) Representative images (left panel) and quantification (right panel) of LOX activity in WT (n=10), *Scx*$^{Cre}$;*Vhl*$^{f/f}$ (n=8) and *Scx*$^{Cre}$;*Vhl*$^{f/f}$;*Hif1α*$^{f/f}$ (n=8) Achilles cryosections. Scale bar: 50μm. (**H, I**) Ramp-to-failure test shows a rescue of E modulus and failure stress in *Scx*$^{Cre}$;*Vhl*$^{f/f}$;*Hif1α*$^{f/f}$ (n=6) TTFs to the WT (n=9) levels compared to *Scx*$^{Cre}$;*Vhl*$^{f/f}$ (n=5) TTFs. n=6 fascicles tested per mouse. (**J, K**) Stress-relaxation test reveals a rescue of stress decay in *Scx*$^{Cre}$;*Vhl*$^{f/f}$;*Hif1α*$^{f/f}$ (n=4) to the WT (n=4) levels compared to *Scx*$^{Cre}$;*Vhl*$^{f/f}$ (n=3) TTFs. n=4 fascicles tested per mouse. (**L**) Representative whole-mount images (left panel) and quantification (right panel) of blood vessels (CD31, red) and myelinated nerves (NM-F, white) in wild type (n=6 for both markers), *Scx*$^{Cre}$;*Vhl*$^{f/f}$ (n=10 for CD31 and n=6 for NM-F marker) and *Scx*$^{Cre}$;*Vhl*$^{f/f}$;*Hif1α*$^{f/f}$ (n=4 for both markers) Achilles tendon. Images show tendon cross-sectional area (YZ plane). White arrows point at the myelinated nerve cells. Scale bar: 100μm. Each point represents a single mouse. Bar graphs indicate mean ± SEM. Kruskal-Wallis with Dunn's multiple comparison test was used in B,D,G,H,I,K,L. Two-way ANOVA with Tukey's multiple comparisons test was used in F,J.

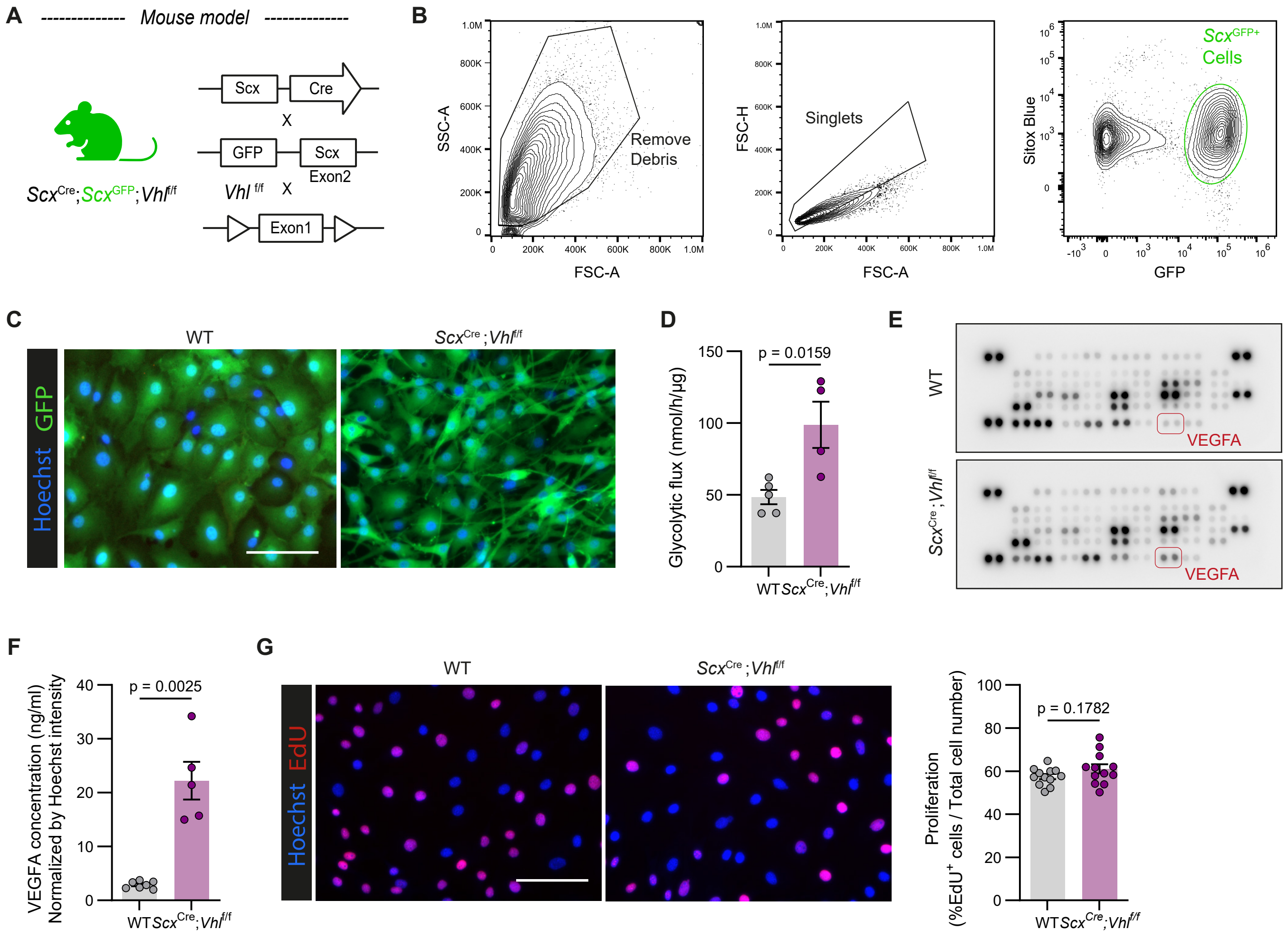
A
Mouse model
Scx
Cre
X
GFP
Scx Exon2
Vhl f/f
X
Exon1
ScxCre;ScxGFP;Vhlf/f
B
SSC-A
FSC-A
Remove Debris
FSC-H
Singlets
Sitox Blue
GFP
ScxGFP+ Cells
C
WT
ScxCre;Vhlf/f
Hoechst GFP
D
Glycolytic flux (nmol/h/µg)
p = 0.0159
WT
ScxCre;Vhlf/f
E
WT
ScxCre;Vhlf/f
VEGFA
F
VEGFA concentration (ng/ml) Normalized by Hoechst intensity
p = 0.0025
G
Hoechst EdU
Proliferation (%EdU+ cells / Total cell number)
p = 0.1782

**Fig. S7. *In vitro* characterization of $Scx^{GFP+}$ tendon cells sorted from TTFs.**

(**A**) Generation of $Scx^{Cre}$; $Scx^{GFP+}$;$Vhl^{f/f}$ reporter mice expressing constitutive Cre and the GFP reporter under the *Scx* promoter. (**B**) Flow cytometry gating strategy for sorting $Scx^{GFP+}$ cells from mouse TTFs. (**C**) Representative images of sorted $Scx^{GFP+}$ cells (Passage 2) from WT and $Scx^{Cre}$; $Scx^{GFP+}$;$Vhl^{f/f}$ TTFs. (**D**) Glycolytic flux in WT and $GFP^{+}$-*Vhl-KO* cells at passage 2 (**E**) Representative images of angiogenic cytokine profile array from WT cells- conditioned media and $GFP^{+}$-*Vhl-KO* cells-conditioned media. (**F**) VEGFA concentration in conditioned media derived from WT and $GFP^{+}$-*Vhl-KO* cells normalized by protein concentration (**G**) Representative images (left panel) and quantification (right panel) of $EdU^{+}$ $Scx^{GFP+}$ cells sorted from WT and $Scx^{Cre}$; $Scx^{GFP+}$;$Vhl^{f/f}$ TTFs. Each dot represents a technical replicate (n=6; 2 technical replicate each). Each point represents a single mouse. Bar graphs indicate mean ± SEM. Student's t test (unpaired, two tailed) was used in D,F,G.

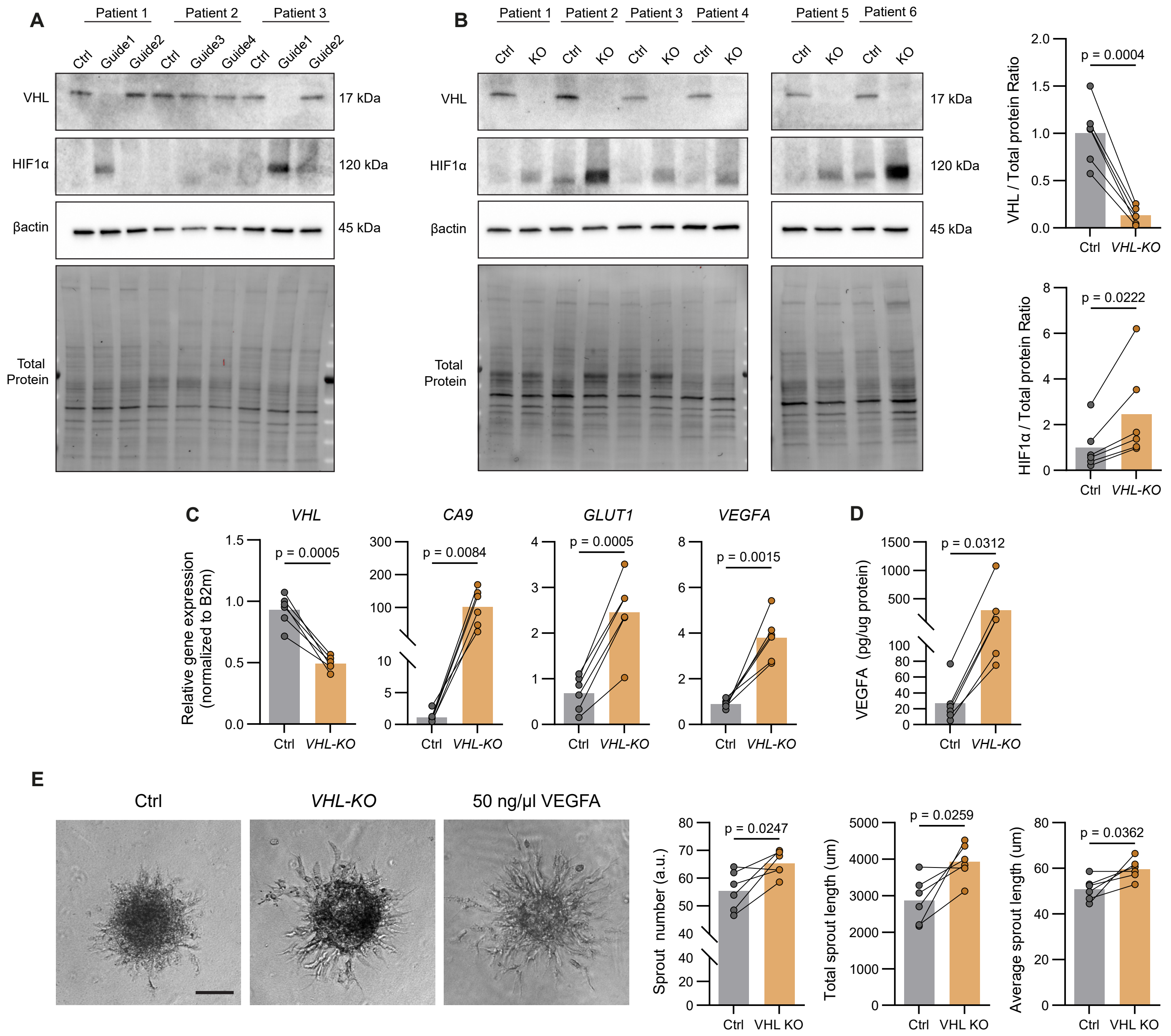

A
Patient 1
Patient 2
Patient 3
Ctrl
Guide1
Guide2
Ctrl
Guide3
Guide4
Ctrl
Guide1
Guide2
VHL
HIF1α
βactin
Total Protein
17 kDa
120 kDa
45 kDa
B
Patient 1
Patient 2
Patient 3
Patient 4
Patient 5
Patient 6
Ctrl
KO
17 kDa
120 kDa
45 kDa
VHL / Total protein Ratio
p = 0.0004
HIF1α / Total protein Ratio
p = 0.0222
Ctrl
VHL-KO
C
VHL
CA9
GLUT1
VEGFA
Relative gene expression (normalized to B2m)
p = 0.0005
p = 0.0084
p = 0.0005
p = 0.0015
D
VEGFA (pg/ug protein)
p = 0.0312
E
Ctrl
VHL-KO
50 ng/µl VEGFA
Sprout number (a.u.)
p = 0.0247
Total sprout length (um)
p = 0.0259
Average sprout length (um)
p = 0.0362
Ctrl
VHL KO

**Fig. S8. Generation of CRISPR/Cas9 -mediated VHL knockout human tendon cells.**

(**A**) Western blot of VHL, HIF1α and βactin protein levels to test the CRISPR/Cas9 VHL-knockout efficiency of four different sgRNAs in human tendon cells. (**B**) Western blot (left panel) and quantification (right panels) of VHL, HIF1α protein levels to verify knockout efficiency of Guide1 in 6 different patients. Protein levels are normalized by total protein. βactin was used as loading control. (**C**) mRNA levels of VHL and canonical HIF1α target genes in Ctrl and *VHL-KO* human tendon cells (n=6). (**D**) VEGFA concentration in conditioned media from Ctrl and *VHL-KO* human tendon cells, measured by ELISA and normalized to protein concentration. (**E**) Representative brightfield images and morphometric quantification of HUVECS spheroid sprout number, total sprout length and average sprout length. Spheroids were either treated with Ctrl- or *VHL-KO*- conditioned media. A third group of spheroids treated with 50ng/µl of VEGFA was included as a technical control. Scale bar: 50µm. Data points represent biologically independent samples. Bar graphs indicate mean ± SEM. Student's t test (paired, two tailed) was used for statistical analysis.

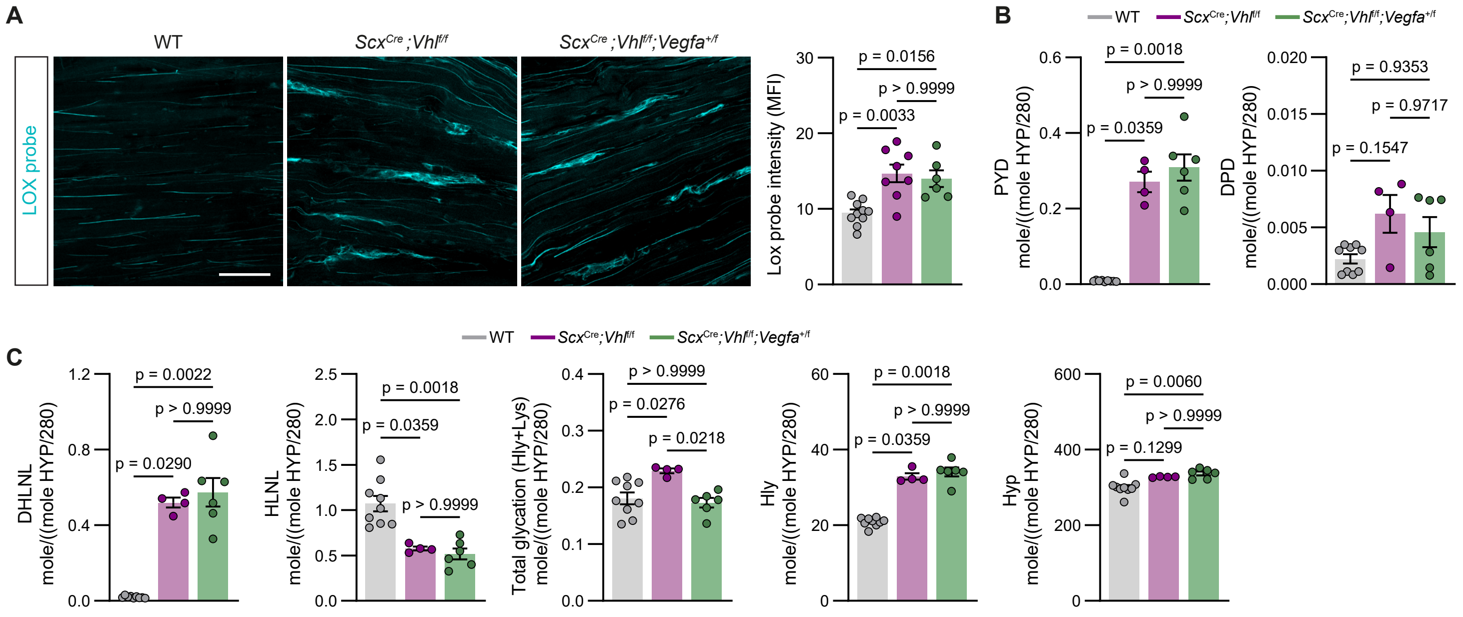
A
WT
ScxCre;Vhlf/f
ScxCre;Vhlf/f;Vegfa+/f
LOX probe
Lox probe intensity (MFI)
p = 0.0156
p > 0.9999
p = 0.0033
B
WT
ScxCre;Vhlf/f
ScxCre;Vhlf/f;Vegfa+/f
PYD
mole/((mole HYP/280)
p = 0.0018
p > 0.9999
p = 0.0359
DPD
mole/((mole HYP/280)
p = 0.9353
p = 0.9717
p = 0.1547
C
DHLNL
mole/((mole HYP/280)
p = 0.0022
p > 0.9999
p = 0.0290
HLNL
mole/((mole HYP/280)
p = 0.0018
p = 0.0359
p > 0.9999
Total glycation (Hly+Lys)
mole/((mole HYP/280)
p > 0.9999
p = 0.0276
p = 0.0218
Hly
mole/((mole HYP/280)
p = 0.0018
p > 0.9999
p = 0.0359
Hyp
mole/((mole HYP/280)
p = 0.0060
p > 0.9999
p = 0.1299

**Fig. S9. *Scx*$^{Cre}$;*Vhl*$^{f/f}$;*Vegfa*$^{+/f}$ mice tendons display levels of LOX activity and collagen crosslinking comparable to *Scx*$^{Cre}$;*Vhl*$^{f/f}$ mice.** (**A**) Representative images (left panel) and quantification (right panel) of LOX activity by utilizing a fluorescent probe in WT (n=10), *Scx*$^{Cre}$;*Vhl*$^{f/f}$ (n=8) and *Scx*$^{Cre}$;*Vhl*$^{f/f}$;*Vegfa*$^{+/f}$ (n=6) Achilles cryosections. Scale bar: 50µm. (**B**) PYD, DPD and (**C**) DHLNL, HLNL, total glycation, hydroxylysine and hydroxyproline content normalized by Hyp content divided by total number of Hyp residues in collagen I (280) in WT (n=9), *Scx*$^{Cre}$;*Vhl*$^{f/f}$ (n=4) and *Scx*$^{Cre}$;*Vhl*$^{f/f}$;*Vegfa*$^{+/f}$ (n=6) TTFs. Each point represents a single mouse. Bar graphs indicate mean ± SEM. Kruskal-Wallis with Dunn's multiple comparison test was used for statistical analysis.

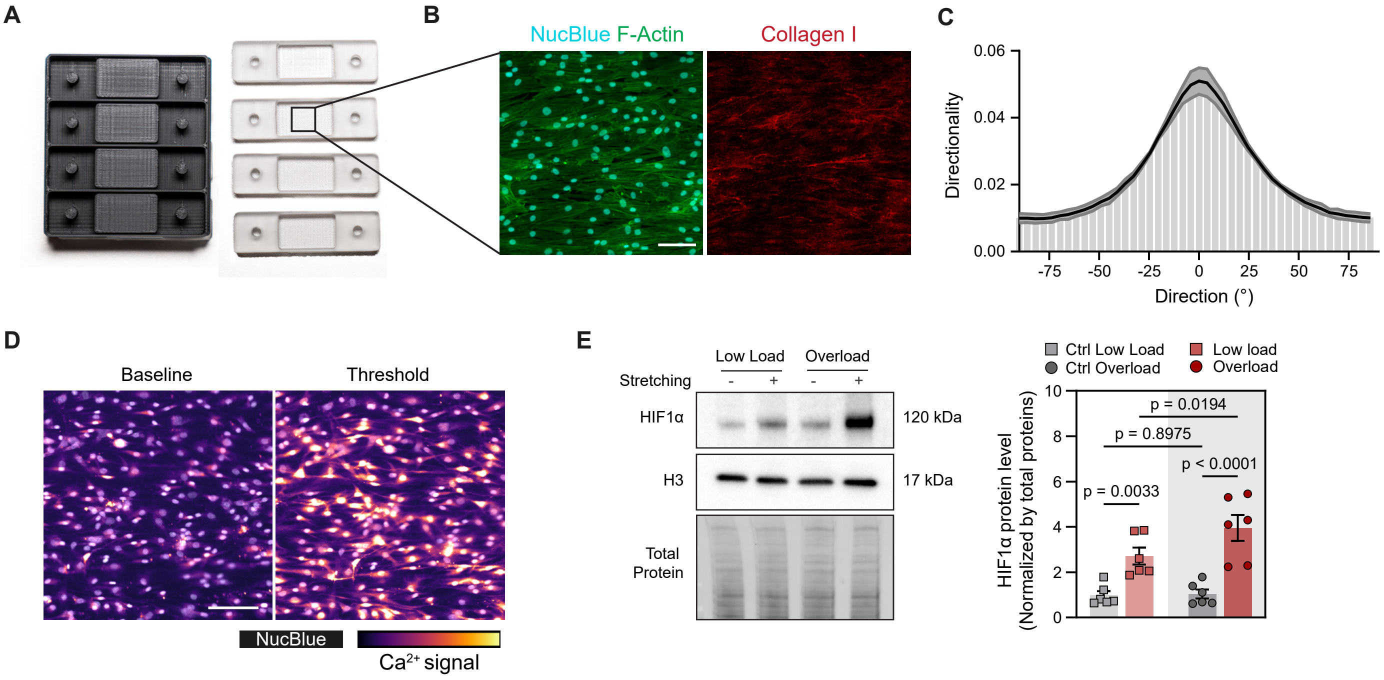
A
B
NucBlue F-Actin
Collagen I
C
Directionality
0.06
0.04
0.02
0.00
-75
-50
-25
0
25
50
75
Direction (°)
D
Baseline
Threshold
NucBlue
Ca2+ signal
E
Low Load
Overload
Stretching
-
+
-
+
HIF1α
120 kDa
H3
17 kDa
Total Protein
Ctrl Low Load
Ctrl Overload
Low load
Overload
HIF1α protein level (Normalized by total proteins)
10
8
6
4
2
0
p = 0.0194
p = 0.8975
p < 0.0001
p = 0.0033

**Fig. S10. Characterization of 2D silicone chambers for cyclic, uniaxial tensile stretching.**

(**A**) Representative images of 3D-printed models and single-use stretchable silicone chambers. (**B**) Representative images of human-derived tendon fibroblasts seeded on the silicone chamber stained for F-actin (Phalloidin, green), collagen I (red) and cell nuclei (NucBlue, cyan). Scale bar: 100μm. (**C**) Cell alignment quantification with ImageJ Directionality plugin. Three biological replicates (n=3 donors) were tested with three technical replicates (n=9 chambers). Black and grey lines indicate mean ± SD. (**D**) Representative $Ca^{2+}$ signaling images of human-derived tendon fibroblasts at baseline and during the stretching protocol showing stretch-induced $Ca^{2+}$ signals. Scale bar: 100μm. (**E**) Quantification of HIF1α protein levels in human tenocytes subjected to low load and overload stretching for 4 hours (n=6). Data points represent biologically independent samples. Bar graphs indicate mean ± SEM. Kruskal-Wallis with Dunn's multiple comparison test was used for statistical analysis in E.

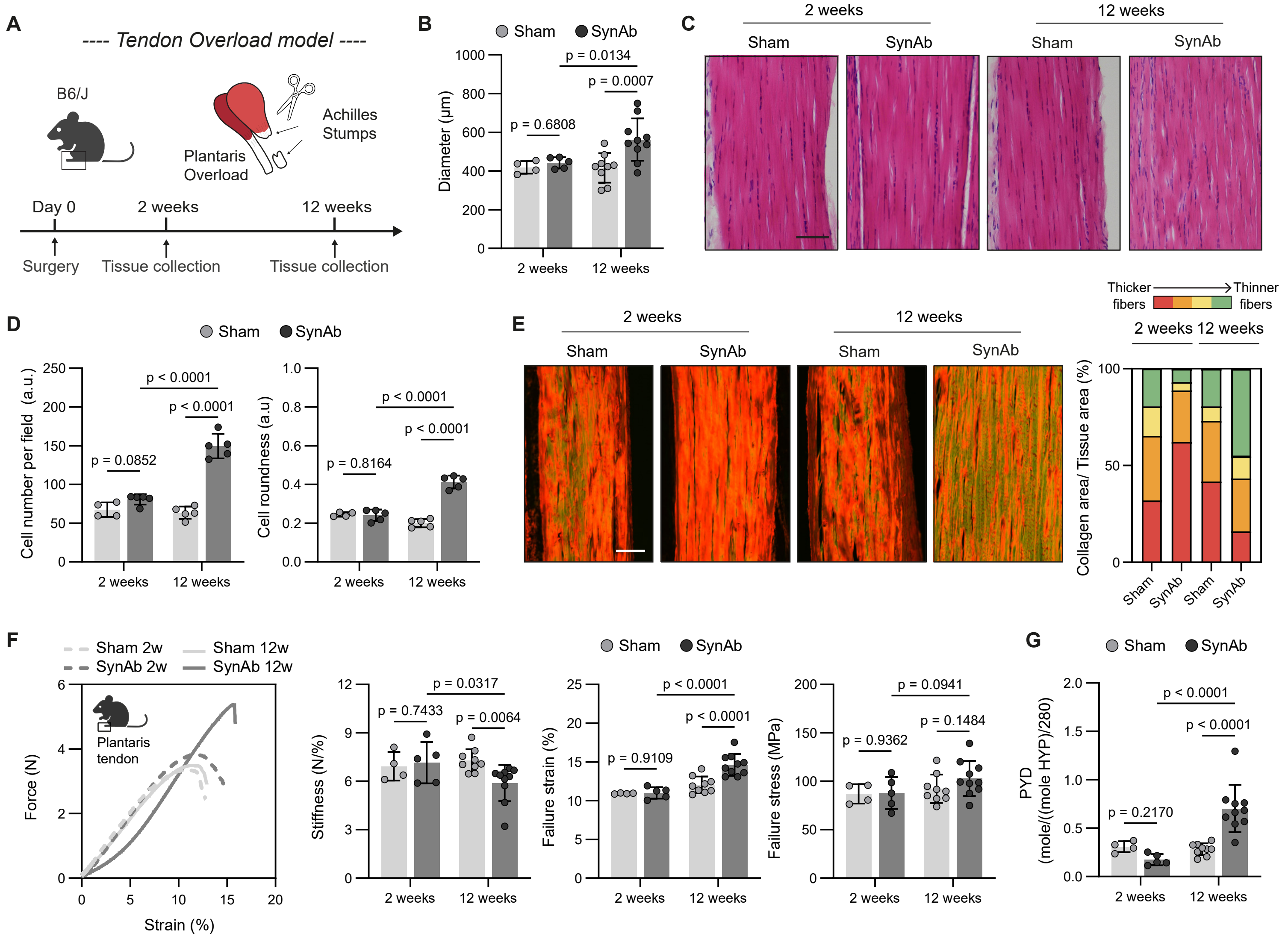
A
---- Tendon Overload model ----
B6/J
Achilles
Stumps
Plantaris
Overload
Day 0
2 weeks
12 weeks
Surgery
Tissue collection
Tissue collection
B
Sham
SynAb
Diameter (μm)
p = 0.0134
p = 0.0007
p = 0.6808
2 weeks
12 weeks
C
2 weeks
12 weeks
Sham
SynAb
Sham
SynAb
D
Sham
SynAb
Cell number per field (a.u.)
p < 0.0001
p < 0.0001
p = 0.0852
Cell roundness (a.u)
p < 0.0001
p < 0.0001
p = 0.8164
2 weeks
12 weeks
E
2 weeks
12 weeks
Sham
SynAb
Sham
SynAb
Thicker fibers
Thinner fibers
2 weeks 12 weeks
Collagen area/ Tissue area (%)
Sham
SynAb
Sham
SynAb
F
Sham 2w
SynAb 2w
Sham 12w
SynAb 12w
Plantaris tendon
Force (N)
Strain (%)
Sham
SynAb
Stiffness (N/%)
p = 0.0317
p = 0.7433
p = 0.0064
Failure strain (%)
p < 0.0001
p < 0.0001
p = 0.9109
Failure stress (MPa)
p = 0.0941
p = 0.1484
p = 0.9362
2 weeks
12 weeks
G
Sham
SynAb
PYD (mole/((mole HYP)/280)
p < 0.0001
p < 0.0001
p = 0.2170
2 weeks
12 weeks

**Fig. S11. Overload induces structural and functional impairments in the plantaris tendon at 12 weeks, but not at 2 weeks.** (**A**) Experimental schematic and timeline of the synergist ablation (SynAb) model performed on B6/J mice. (**B**) Quantification of plantaris tendon diameter at 2 weeks (n=4,Sham; n=5,SynAb) and 12 weeks post-surgery (n=9,Sham; n=10,SynAb). (**C**) Representative images of haematoxylin-eosin (H&E)-stained sections and (**D**) cell number and cell roundness quantification of 2 weeks-overload (n=4,Sham; n=5,SynAb) and 12 weeks-overload (n=9,Sham; n=10,SynAb) plantaris tendon. Scale bar: 50μm. (**E**) Representative images of picrosirius red (PSR) staining under polarized light and qualitative quantification of collagen fibers content in plantaris tendon after 2 weeks (n=4,Sham; n=5,SynAb) and 12 weeks (n=5,Sham; n=5,SynAb) overload. Red, orange, yellow and green colors represent different types of collagen fibers based on their thickness. Red indicates thicker/mature collagen fibers while green indicates thinner/immature fibers. Scale bar: 50μm. (**F**) Ramp-to-failure test shows a decreased stiffness and increased failure strain in plantaris tendon at 12 weeks (n=9,Sham; n=10,SynAb) overload but not at 2 weeks (n=4,Sham; n=5,SynAb). (**G**) Mature PYD crosslinks levels normalized by Hyp content divided by total number of Hyp residues in collagen I (280) of 2 weeks-overload (n=4,Sham; n=5,SynAb) and 12 weeks-overload (n=9,Sham; n=10,SynAb) plantaris tendon. Data points represent biologically independent samples. Bar graphs indicate mean ± SEM. Two-way ANOVA with Tukey's multiple comparisons test was used in B,D, F,G.

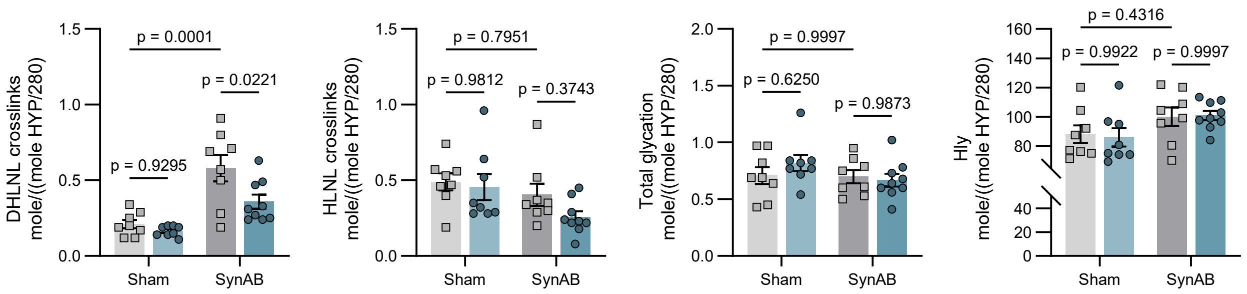
DHLNL crosslinks
mole/((mole HYP/280)
p = 0.0001
p = 0.0221
p = 0.9295
Sham
SynAB
HLNL crosslinks
mole/((mole HYP/280)
p = 0.7951
p = 0.9812
p = 0.3743
Total glycation
mole/((mole HYP/280)
p = 0.9997
p = 0.6250
p = 0.9873
Hly
mole/((mole HYP/280)
p = 0.4316
p = 0.9922
p = 0.9997

**Fig. S12. Overload-induced changes in collagen crosslinks and hydroxyproline levels of plantaris tendon in WT and *Scx*$^{Cre}$;*Hif1α*$^{f/f}$ mice.** DHLNL, HLNL, total glycation and hydroxylysine content normalized by Hyp content divided by total number of Hyp residues in collagen I (280) in WT (n=8 for both conditions) and *Scx*$^{Cre}$;*Vhl*$^{f/f}$ (n=4) and *Scx*$^{Cre}$;*Hif1α*$^{f/f}$ groups (n=8 Sham; n=9 SynAb). Each point represents a single mouse. Bar graphs indicate mean ± SEM. Two-way ANOVA with Tukey's multiple comparisons test was used for statistical analysis.

**Table S1. Patient demographics, surgical indications and tissue analysis techniques.**

| Recorded ID | Tendon Type | Gender | Age | Diabetic | Surgery Reason | Tissue collection | Transcriptomics | Label-free Proteomics | Pathological evaluation (Bonar Score) | HIF1a staining | IF | Crosslinks/ Quantitative Proteomics |
|---|---|---|---|---|---|---|---|---|---|---|---|---|
| 1315 | Long Head of Biceps Tendon | m | 48 | n | Subscapularis tendon lesion | Biceps tenodesis | No | No | Yes | Yes | Yes | No |
| 1317 | Long Head of Biceps Tendon | m | 44 | n | Supraspinatus tendon lesion | Biceps tenodesis | No | No | Yes | Yes | Yes | Yes |
| 1318 | Long Head of Biceps Tendon | m | 35 | n | Type II SLAP lesion | Biceps tenodesis | Yes | No | Yes | Yes | Yes | Yes |
| 1326 | Long Head of Biceps Tendon | m | 73 | n | Rotator cuff rupture | Biceps tenodesis | No | No | Yes | Yes | Yes | Yes |
| 1331 | Long Head Biceps Tendon | f | 37 | n | Type II SLAP lesion and partial articular-sided small supraspinatus lesion | Biceps tenodesis | Yes | No | Yes | Yes | Yes | Yes |
| 1375 | Long Head of Biceps Tendon | m | 56 | n | Subacromial bursitis with incipient Omarthrosis | Biceps tenodesis | Yes | No | Yes | Yes | Yes | Yes |
| 1383 | Long Head of Biceps Tendon | f | 54 | n | SLAP lesion and rotator cuff rupture (supraspinatus transmural, subscapularis upper edge) | Biceps tenodesis | No | No | Yes | Yes | Yes | Yes |
| 1387 | Long Head Biceps Tendon | m | 59 | n | Rotator cuff rupture (supraspinatus and infraspinatus) | Biceps tenodesis | No | No | Yes | Yes | Yes | Yes |
| 1392 | Long Head of Biceps Tendon | m | 48 | n | Rotator cuff rupture (supraspinatus total transmural, infraspinatus anterior half partial) | Biceps tenodesis | No | No | Yes | Yes | Yes | Yes |
| 1399 | Long Head of Biceps Tendon | m | 43 | n | AC joint arthrosis and long head biceps tendinopathy | Biceps tenodesis | Yes | No | Yes | Yes | Yes | Yes |

| | | | | | | | | | | | | |
|---|---|---|---|---|---|---|---|---|---|---|---|---|
| 1400 | Long Head of Biceps Tendon | m | 66 | n | Partial articular-sided rotator cuff lesion (supraspinatus and infraspinatus tendons) | Biceps tenodesis | Yes | No | Yes | Yes | Yes | Yes |
| 1402 | Long Head of Biceps Tendon | m | 36 | n | Type II SLAP lesion | Biceps tenodesis | No | No | Yes | Yes | Yes | Yes |
| 1403 | Long Head of Biceps Tendon | m | 50 | n | Rotator cuff rupture (supraspinatus total transmural, infraspinatus anterior half partial, subscapularis upper third tendons) | Biceps tenodesis | Yes | No | Yes | Yes | Yes | Yes |
| 1404 | Long Head of Biceps Tendon | m | 34 | n | Type II SLAP Lesion | Biceps tenodesis | Yes | No | Yes | Yes | Yes | Yes |
| 1408 | Long Head of Biceps Tendon | m | 32 | n | Type II SLAP lesion | Biceps tenodesis | Yes | No | Yes | Yes | Yes | Yes |
| 1409 | Long Head of Biceps Tendon | m | 46 | n | PASTA lesion | Biceps tenodesis | Yes | No | Yes | Yes | Yes | Yes |
| 1410 | Long Head of Biceps Tendon | m | 40 | n | Type II SLAP lesion | Biceps tenodesis | Yes | No | Yes | Yes | Yes | Yes |
| 1413 | Long Head of Biceps Tendon | m | 57 | n | Supraspinatus partial rupture on the articular side and lesion of the lateral pulley | Biceps tenodesis | No | No | Yes | Yes | Yes | Yes |
| 1418 | Long Head of Biceps Tendon | m | 58 | n | Partial rotator cuff rupture (supraspinatus and subscapularis tendon) | Biceps tenodesis | No | No | Yes | Yes | Yes | Yes |
| 1439 | Long Head of Biceps Tendon | m | 54 | n | Tendinitis Calcarea | Biceps tenodesis | No | No | Yes | Yes | Yes | Yes |
| 1441 | Long Head of Biceps Tendon | m | 42 | n | Supraspinatus subtotal rupture with SLAP lesion | Biceps tenodesis | Yes | No | Yes | Yes | Yes | Yes |
| 1442 | Long Head of Biceps Tendon | m | 46 | n | Painful intra-articular biceps tenodesis from postoperative frozen shoulder arthroscopy | Biceps tenodesis | No | No | Yes | Yes | Yes | Yes |

| | | | | | | | | | | | | |
|---|---|---|---|---|---|---|---|---|---|---|---|---|
| 1446 | Long Head of Biceps Tendon | f | 54 | n | SLAP lesion | Biceps tenodesis | Yes | No | Yes | Yes | Yes | Yes |
| 1448 | Long Head of Biceps Tendon | m | 54 | n | Rotator cuff rupture (supraspinatus) | Biceps tenodesis | No | No | Yes | Yes | Yes | No |
| 1450 | Long Head of Biceps Tendon | m | 47 | n | Rotator cuff rupture (anterior supraspinatus, partial rupture of the subscapularis) | Biceps tenodesis | No | No | Yes | Yes | Yes | Yes |
| 1453 | Long Head of Biceps Tendon | m | 52 | n | PASTA lesion, SLAP lesion | Biceps tenodesis | Yes | No | Yes | Yes | Yes | Yes |
| 1456 | Long Head of Biceps Tendon | m | 34 | n | Subluxation of the long head of Biceps tendon with upper edge lesion of subscapularis tendon | Biceps tenodesis | Yes | No | Yes | Yes | Yes | Yes |
| 1459 | Long Head of Biceps Tendon | f | 55 | n | SLAP lesion | Biceps tenodesis | Yes | No | Yes | Yes | Yes | Yes |
| 1061 | Gracilis Tendon | m | 51 | n | ACL rupture | ACL reconstruction / graft tissue | No | No | Yes | Yes | Yes | Yes |
| 1066 | Gracilis Tendon | f | 45 | n | Patellar instability | MPFL reconstruction / graft tissue | No | No | Yes | Yes | Yes | Yes |
| 1095 | Gracilis Tendon | m | 17 | n | ACL rupture | ACL reconstruction / graft tissue | No | No | Yes | Yes | Yes | Yes |
| 1097 | Gracilis Tendon | f | 21 | n | ACL rupture | ACL reconstruction / graft tissue | No | No | Yes | Yes | Yes | Yes |
| 1111 | Gracilis Tendon | m | 28 | n | ACL rupture | ACL reconstruction / graft tissue | No | No | Yes | Yes | Yes | Yes |
| 1143 | Gracilis Tendon | m | 40 | n | Patellar instability | MPFL reconstruction / graft tissue | No | No | Yes | Yes | Yes | Yes |
| 1156 | Gracilis Tendon | f | 45 | n | ACL rupture | ACL reconstruction / graft tissue | No | No | Yes | Yes | Yes | Yes |

| | | | | | | | | | | | | |
|---|---|---|---|---|---|---|---|---|---|---|---|---|
| 1165 | Gracilis Tendon | m | 25 | n | ACL rupture | ACL reconstruction / graft tissue | No | No | Yes | Yes | Yes | Yes |
| 1200 | Gracilis Tendon | m | 21 | n | Patellar instability | MPFL reconstruction / graft tissue | No | No | Yes | Yes | Yes | Yes |
| 1202 | Gracilis Tendon | f | 36 | n | Patellar instability | MPFL reconstruction / graft tissue | No | No | Yes | Yes | Yes | Yes |
| 1210 | Gracilis Tendon | f | 15 | n | MPFL rupture | ACL reconstruction / graft tissue | No | No | Yes | Yes | Yes | Yes |
| 1221 | Gracilis Tendon | m | 28 | n | ACL rupture | ACL reconstruction / graft tissue | No | No | Yes | Yes | Yes | Yes |
| 1227 | Gracilis Tendon | f | 31 | n | ACL rupture | ACL reconstruction / graft tissue | No | No | Yes | Yes | Yes | Yes |
| 1232 | Gracilis Tendon | f | 20 | n | Patellofemoral instability | MPFL reconstruction / graft tissue | No | No | Yes | Yes | Yes | Yes |
| 1234 | Gracilis Tendon | f | 33 | n | ACL rupture | ACL reconstruction / graft tissue | No | No | Yes | Yes | Yes | Yes |
| 1248 | Gracilis Tendon | m | 41 | n | ACL rupture | ACL reconstruction / graft tissue | No | No | Yes | Yes | Yes | Yes |
| 1381 | Gracilis Tendon | f | 40 | n | ACL rupture | ACL reconstruction / graft tissue | No | No | Yes | Yes | Yes | Yes |
| 853a | Gracilis Tendon | f | 52 | n | ACL rupture | ACL reconstruction / graft tissue | No | Yes | No | No | No | No |
| 207b | Gracilis Tendon | m | 59 | n | ACL rupture | ACL reconstruction / graft tissue | No | Yes | No | No | No | No |
| 284a | Gracilis Tendon | f | 50 | n | ACL rupture | ACL reconstruction / graft tissue | No | Yes | No | No | No | No |

| | | | | | | | | | | | |
|---|---|---|---|---|---|---|---|---|---|---|---|
| 852b | Gracilis Tendon | f | 48 | n | ACL rupture | ACL reconstruction / graft tissue | No | Yes | No | No | No | No |
| 736110 | Rotator cuff tendon | f | 65 | n | Rotator cuff rupture (supraspinatus/infraspinatus transmural, subscapularis 50%) | Supraspinatus Biopsy | No | Yes | No | No | No | No |
| 793421 | Rotator cuff tendon | m | 65 | n | Traumatic Rotator cuff rupture (supraspinatus, infraspinatus, subscapularis upper edge) | Supraspinatus Biopsy | No | Yes | No | No | No | No |
| 750464 | Rotator cuff tendon | f | 45 | n | Traumatic rotator cuff rupture (supraspinatus transmural, subscapularis upper edge) | Supraspinatus Biopsy | No | Yes | No | No | No | No |
| 308629 | Rotator cuff tendon | m | 67 | n | Traumatic rotator cuff rupture (supraspinatus and subscapularis, partial infraspinatus) | Supraspinatus Biopsy | No | Yes | No | No | No | No |

**Table S2. Mouse lines information.**

| Mouse Alleles | | | | | |
|---|---|---|---|---|---|
| **Mouse allele cited in text** | **Allele full name** | **Supplier** | **Stock number** | **Citation** | **Pubmed ID** |
| $Scx^{Cre}$ | tg(Scx-GFP/cre)1Stzr | Gift from Dr. Ronen Schweitzer (OSHU) | N/A | Blitz et al. (2009) Dev Cell. 17(6):861-73. | 20059955 |
| $Scx^{GFP}$ | tg(ScxGFP) | Gift from Dr. Ronen Schweitzer (OSHU) | N/A | Pryce et al. (2007) Dev Dyn 236(6):1677-82. | 17497702 |
| $Hif1\alpha^{fl/fl}$ | B6.129-Hif1atm3Rsjo/J | JAX | 007561 | Ryan et al. (2000) Cancer Res. 60(15):4010-5. | 10945599 |
| $Vegfa^{fl/fl}$ | B6.Cg-VEGF flox/flox | Gift from Christian Stockmann (Zurich) | N/A | Stockmann et al. (2008) 456(7223):814-8. | 18997773 |
| $Vhl^{fl/fl}$ | B6.Cg-VHL flox/flox | Gift from Christian Stockmann (Zurich) | N/A | Haase at al. (2001) Proc Natl Acad Sci USA. 98(4):1583-8 | 11171994 |
| B6/J | C57BL/6J | JAX | 000664 | Simon MM et al. (2013) Genome Biol. 14(7):R82. | 23902802 |

**Table S3. Sequences of single guide RNA used for CRISPR/Cas9 -mediated VHL knockout of human tendon cells. Related to Figure S8.**

| Guide name | Sequence |
|---|---|
| sgRNA1_VHL | CGCGCGTCGTGCTGCCCGTA |
| sgRNA2_VHL | TCTCTCAATGTTGACGGACA |
| sgRNA3_VHL | GGTCATCTTCTGCAATCGCA |
| sgRNA4_VHL | ATGGATTCATGGAGTAGCCT |
| sgRNA_AAVS1 | GGGGCCACTAGGGACAGGAT |

**Table S4. Antibodies List**

| Antibodies used for immunofluorescence | | |
|---|---|---|
| ***Primary antibodies*** | ***Source*** | ***Identifier*** |
| Goat anti-CD31/PECAM1 | R&D Systems | AF3628 |
| Rabbit anti-Neurofilament M (NF-M) | Biolegend | 841001 |
| Rabbit anti-HIF1α | Cell signaling Technology | 36169 |
| CellMask Deep Red Plasma Membrane Stain | Thermo Fisher Scientific | C10046 |
| Alexa Fluor® 488 Phalloidin | Thermo Fisher Scientific | A12379 |
| Alexa Fluor® 568 Phalloidin | Thermo Fisher Scientific | A12380 |
| Rabbit anti-Collagen I | Boster | PA2140-2 |
| Fluo-4, AM, cell permeant | Thermo Fisher Scientific | F14217 |
| ***Secondary antibodies*** | ***Source*** | ***Identifier*** |
| Alexa Fluor 647 anti-Goat | Thermo Fisher Scientific | A21447 |
| Alexa Fluor 546 anti-Rabbit | Thermo Fisher Scientific | A11010 |
| Alexa Fluor 568 anti-Rabbit | Thermo Fisher Scientific | A10042 |
| **Antibodies used for Immunoblot** | | |
| ***Primary antibodies*** | ***Source*** | ***Identifier*** |
| Rabbit anti-HIF1α | Cell signaling Technology | 36169 |
| Rabbit anti-HIF1α | Cayman Chemical | 10006421 |
| Rabbit anti-HIF2α | Novus Biological | NB100-122SS |
| Rabbit anti-VHL | Cell signaling | 68547S |
| Rabbit anti-VHL | GeneTex | GTX101087 |
| Mouse anti-Histone H3 | Cell signaling Technology | 14269S |
| Mouse anti-β-Actin | Cell signaling Technology | 3700T |
| ***Secondary antibodies*** | ***Source*** | ***Identifier*** |
| Anti-Rabbit IgG (H+L), highly cross adsorbed-Peroxidase antibody produced in goat | Sigma-Aldrich | SAB3700878 |
| Anti-Mouse IgG (H+L), highly cross adsorbed-Peroxidase antibody produced in goat | Sigma-Aldrich | SAB3701073 |
| **Antibodies used for flow cytometry** | | |
| PE Rat Anti-Mouse CD31 [MEC 13.3] | BD Biosciences | 553373 |
| APC/fire™ 750 anti-mouse CD45 [30-F11] | Biolegend | 103154 |
| SYTOX® Blue Dead Cell Stain | Thermo Fisher Scientific | S34857 |

**Table S5. Sequences of primers used for RT-PCR. Related to Figure S3 and Figure S8.**

| Primer name | Species | Forward sequence | Reverse sequence |
|---|---|---|---|
| *Vhl* | *Mus Musculus* | ATCCACAGCTACCGAGGTCA | TCGACATTGAGGGATGGCAC |
| *Car9* | *Mus Musculus* | GCGCTAAGCAGCTCCATACTC | CGTGGCTCGGAAGTTCAGTT |
| *Vegfa* | *Mus Musculus* | CTGTACCTCCACCATGCCAAGT | TCGCTGGTAGACATCCATGAACT |
| *Glut1* | *Mus Musculus* | GTGGTGAGTGTGGTGGATG | AGTTCGGCTATAACACTGGTG |
| *Phd3* | *Mus Musculus* | CAGACCGCAGGAATCCACAT | TTCAGCATCGAAGTACCAGAC |
| *Slc7a5* | *Mus Musculus* | CTGGTCTTCGCCACCTACTT | GCCTTTACGCTGTAGCAGTTC |
| *Anxa5* | *Mus Musculus* | AGCATCATGGCTACGAGAGG | GCTTCGGGATGTACCCAGGT |
| *B2m* | *Mus Musculus* | CACTGAATTCACCCCCACTGA | CGATCCCAGTAGACGGTCTTG |
| **Primer name** | **Species** | **TaqMan code (IDT)** | |
| *VHL* | *Homo Sapiens* | Hs.PT.58.24810691.g | |
| *CA9* | *Homo Sapiens* | Hs.PT.56a.2192890 | |
| *GLUT1* | *Homo Sapiens* | Hs.PT.58.25872862 | |
| *VEGFA* | *Homo Sapiens* | Hs.PT.58.1149801 | |
| *ANXA5* | *Homo Sapiens* | Hs.PT.56a.38876508 | |
| *B2M* | *Homo Sapiens* | Hs.PT.58v.18759587 | |
